# Targeted fluorescent lipid microparticles for quantitative measurement of phagosomal pH

**DOI:** 10.1101/2025.11.20.685382

**Authors:** Sophie Michelis, Héloïse Uhl, Florence Niedergang, Jacques Fattaccioli, Blaise Dumat, Jean-Maurice Mallet

## Abstract

Phagosomal acidification plays a pivotal role in pathogen destruction and immune signaling, yet tools capable of reporting these biochemical changes with spatial and mechanistic precision remain scarce. A modular biosensing strategy is presented in which BODIPY-derived hydrophilic fluorophores are conjugated to phospholipids and incorporated at the surface of targeted lipid microparticles. These soft and biomimetic particles combine receptor-specific uptake with ratiometric fluorescence readouts, enabled by pairing pH-responsive dyes at the interface with a pH-invariant reference probe in the core. Following Fcγ receptor-mediated internalization by macrophages, the particles deliver real-time insight into the onset and progression of phagosomal acidification. This versatile platform provides a direct means to couple defined surface recognition events to intracellular pH measurements, offering new opportunities to unravel how particle identity and ligand presentation modulate phagosomal physiology.

## INTRODUCTION

Phagocytosis is a receptor-mediated internalization and digestion process of objects larger than 0.5 microns leading to the formation of a phagosome. It is accompanied by dynamic variations of the phagosomal pH, which play a central role in regulating degradation, antigen processing, and immune signaling.^[1,2]^ After internalization, phagosomes undergo progressive acidification driven by the recruitment of vacuolar H^+^-ATPases (V-ATPases) and the fusion with endosomal and lysosomal compartments.^[3]^ The resulting decrease in pH activates hydrolytic enzymes and provides an optimal environment for pathogen killing and antigen presentation. Monitoring these changes requires probes that are efficiently internalized by phagocytic cells and that report local acidification with high specificity.

Several fluorescence microscopy–based approaches have been developed to monitor phagosomal maturation and acidification. Freely-diffusible small molecular probes for pH or enzymatic can be used but, in the absence of conjugation to the phagocytosed object, they provide a global view of phagosomal activation dynamics rather than tracking individual internalization events.^[4,5]^ To achieve more detailed measurements, pH-sensitive conjugates of bacteria or silica microparticles —typically using commercially-available probes such as fluorescein isothiocyanate (FITC) or pHrodo red/green— have been employed. These particles are efficiently internalized by macrophages and are compatible with both flow cytometry and fluorescence microscopy analyses.^[3,6–11]^ Early work using FITC-labeled *S. aureus* revealed a rapid loss of fluorescence consistent with phagosomal acidification,^[3]^ while studies with FITC-labeled silica beads reported similar changes occurring with distinct kinetics.^[7]^ In more recent results, IgG-coated silica beads incorporating pHrodo red provided evidence for a multistage acidification trajectory, involving a lag phase followed by acidification.^[9]^ Using Janus particles, the same research group further demonstrated that the spatial presentation of ligands on the particle surface modulates the kinetics of phagosomal maturation.^[12]^

While bacterial conjugates remain valuable tools, they activate a complex and heterogeneous set of membrane receptors. In contrast, solid silica particles enable more selective engagement of receptors such as Fcγ receptors or dectin-1, although their mechanical properties differ from that of biological particles such as bacteria.^[12]^ Collectively, these studies highlight that phagosomal maturation is influenced by the physicochemical nature of the target particle and the molecular cues it presents and also —as shown previously— by the type of receptor engaged.^[13,14]^

To study individual phagocytic events and follow phagosome maturation, we developed a biomimetic ratiometric pH sensor for monitoring phagosomal acidification, based on targeted lipid microparticles.^[15–18]^

To this end, we synthesized a family of water-soluble, bioconjugatable BODIPY-derived pH probes (**BODIpH**) with emission wavelengths spanning from yellow to far-red. These probes display a turn-on fluorescence from neutral to acidic pH with high dynamic range and p*K*_*a*_ values that can be tuned from 8 to 6, making them well suited for monitoring pH variations during endocytic processes (Figure 1A). Following conjugation to phospholipids, these probes were incorporated onto the surface of micrometer-sized oil-in-water droplets, together with biotinylated ligands allowing opsonization with anti-biotin immunoglobulin G (IgG) (Figure 1B).^[15,17]^ In contrast to solid silica particles, the fluid interface allows the phagocytic ligands to laterally cluster, better reproducing cell–cell interactions and enabling efficient recognition even by low-affinity ligands such as monosaccharides that target lectin receptors.^[15,16]^A pH-agnostic lipophilic dye was also encapsulated inside the droplets to provide an internal reference for ratiometric quantification of the phagosomal pH. The design thus combines controlled surface functionalization with targeting ligands to trigger phagocytosis and fluorescent pH probes and the resulting lipid microparticles were phagocytosed by RAW 264.7 macrophages through Fcγ receptors engagement, enabling the measurement of phagosomal acidification over time during the maturation stage (Figure 1C&D).

**Figure 1.**
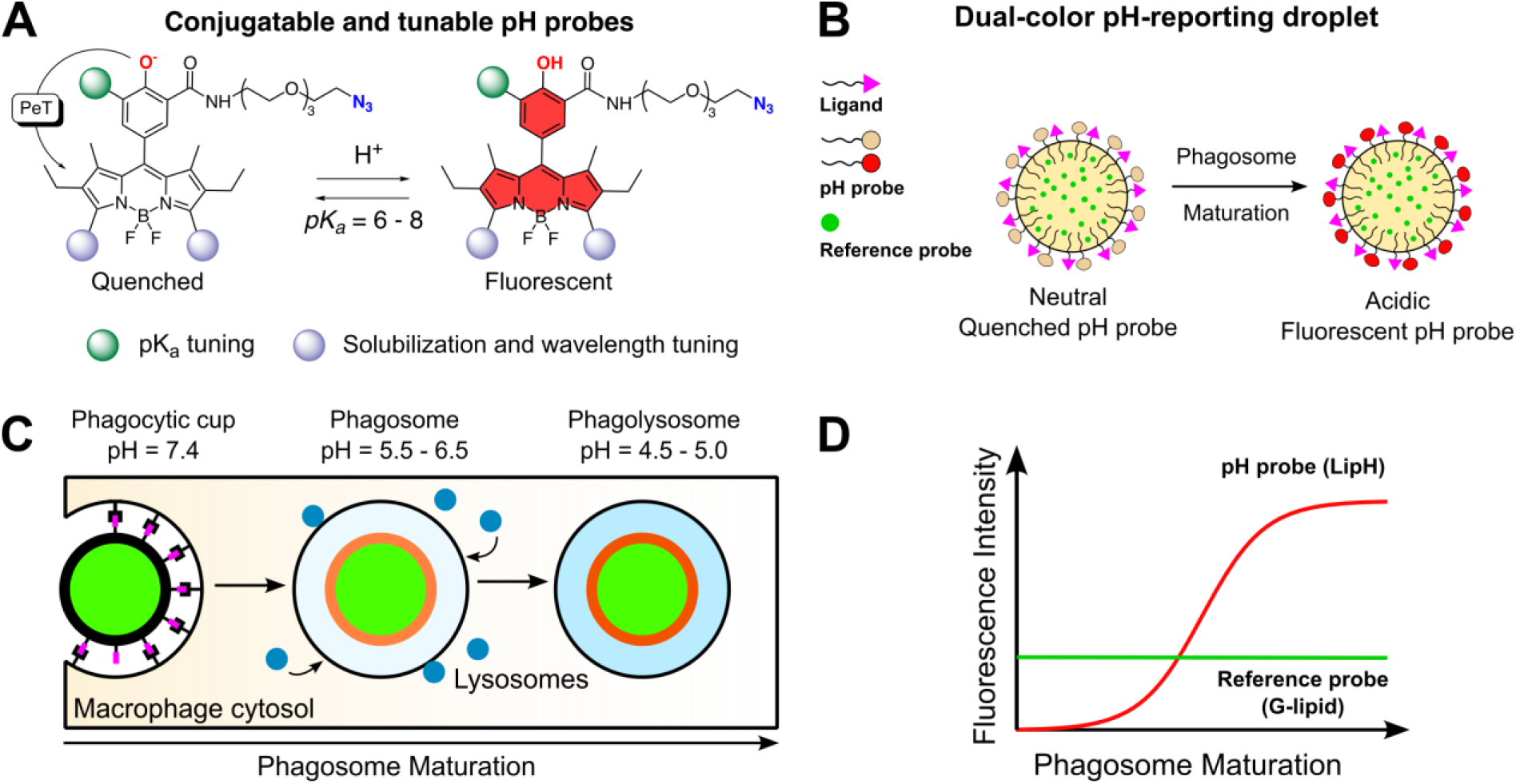
Schematic overview of the design and application of ratiometric pH-sensitive lipid microparticles for monitoring phagosomal acidification. (A) **Design of hydrosoluble pH probes (BODIpH):** BODIPY-derived fluorophores are rendered pH-sensitive by a phenol group that quenches fluorescence through PeT at neutral pH and restores emission upon protonation. An azido linker allows bioconjugation and p*K*_*a*_ and emission wavelength can be tuned. (B) **Lipid microparticle formulation:** BODIpH probes are conjugated to phospholipids and incorporated at the droplet surface, together with a pH-insensitive G-lipid reference and targeting ligands for receptor-mediated uptake. (C) **Quantitative pH sensing during phagocytosis:** Following receptor engagement and internalization by macrophages, droplets sequentially encounter phagosomal environments with decreasing pH (7.4 → 5.5–6.5 → 4.5–5). (D) Ratiometric fluorescence readouts enable quantitative measurement of phagosomal acidification dynamics.

## RESULTS AND DISCUSSION

### BODIpH probes: hydrophilic pH probes with tunable emission wavelengths

Given the central physiological role of pH, a wide range of fluorescent probes has been developed for its measurement. Most of them are based on rather lipophilic neutral or cationic probes that readily permeate the plasma membrane and enable intracellular pH sensing. Following conjugation to biomolecules or particles such lipophilic probes can lead to aggregation and quenching, which can be harnessed as a useful sensing mechanism.^[19,20]^ However, in the frame of this project it would interfere with environmental pH sensing and hydrosoluble probes are thus necessary. Phagocytosis imaging assays frequently require multiplexed fluorescence experiments to visualize additional cellular components —such as actin, receptors, or endosomal markers— alongside exogenous pH sensors. Extending the spectral range of pH-sensitive probes toward the far-red region is therefore essential to enable efficient spectral separation and multicolor imaging. Considering these requirements, commonly used probes for phagocytic assays, such as fluorescein (FITC) or pHrodo red and green, remain suboptimal: fluorescein exhibits a turn-off response and low emission wavelength, while pHrodo dyes display p*K*_*a*_ values around 6.8, resulting in residual fluorescence at neutral pH and a reduced signal-to-noise ratio. To address these limitations, we set out to design new hydrosoluble pH probes with tunable optical properties and p*K*_*a*_ values, suitable for particle surface functionalization and multiplexed monitoring of phagosomal acidification.

The core scaffold of the BODIpH is an ethyl-BODIPY structure (**Yellow BODIpH**) (Figure 2, Table 1). The pH sensitivity stems from a photoinduced electron transfer (PeT) with a phenol moiety in the meso position of the BODIPY: PeT between the phenolate and the fluorophore quenches the fluorescence that is restored upon protonation creating an on/off emission trigger actuated by protons (**Figure 1A**).^[21,22]^ An amide function in ortho position of the phenol reduces the p*K*_a_ of the phenol by approximately two pH units (pKa ≈ 8, BODIpH series 1) thanks to hydrogen bonding.^[23]^ It was also used to introduce a conjugatable handle containing an azide allowing bioconjugation. Addition of an electron-withdrawing bromine atom further decreases the pKa down to around 6 (BODIpH series 2). The synthetic procedures are presented and discussed in the supplementary material (**Scheme S1** and **S2, Table S1**).

**Figure 2.**
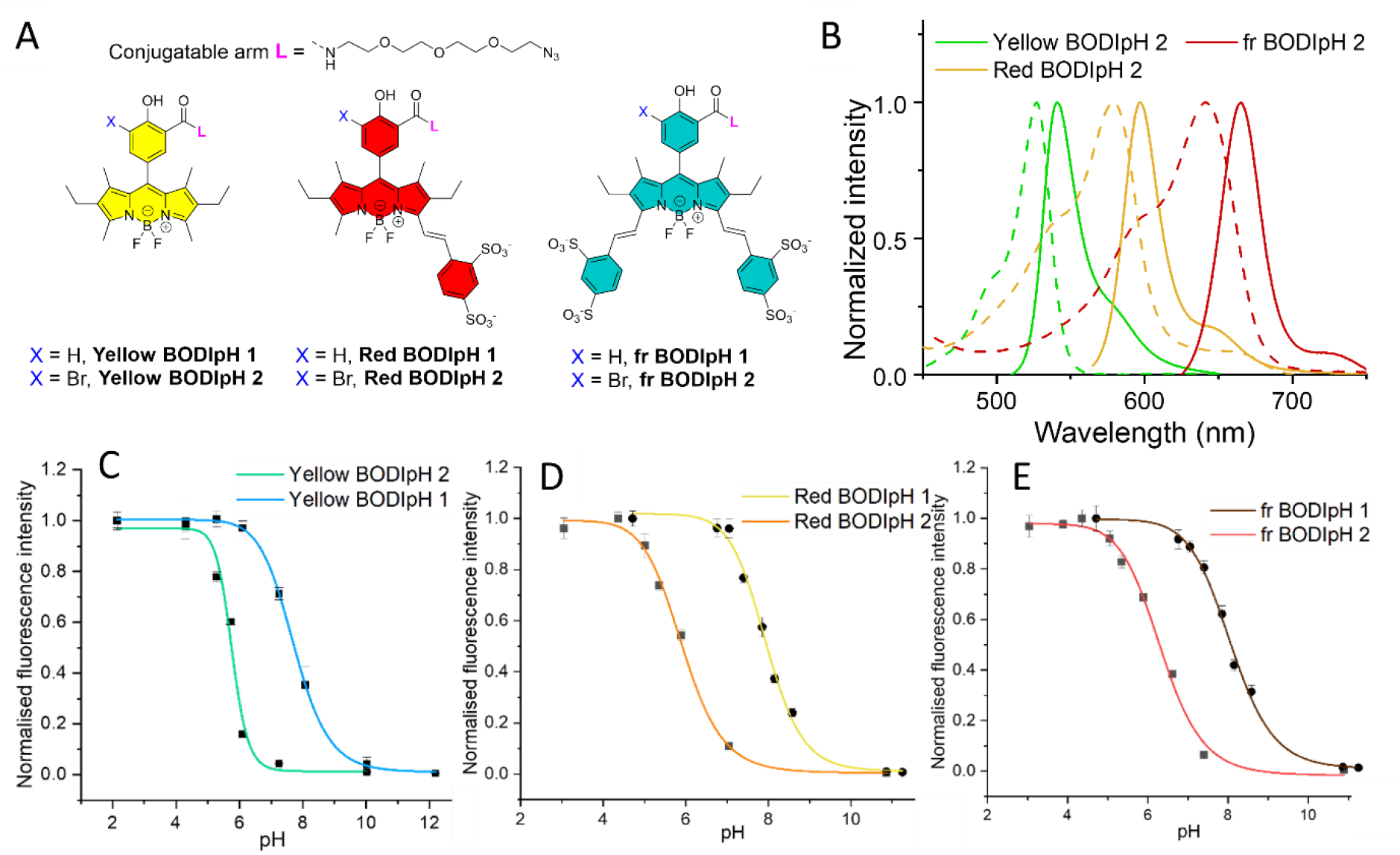
Design and photophysical properties of BODIpH probes. (A) Chemical structures of the six BODIpH derivatives. (B) Normalized absorption (dashed lines) and emission (solid lines) spectra. Fluorometric pH titration curves for (C) Yellow BODIpH, (D) Red BODIpH, and (E) far-red (fr) BODIpH. Titrations were performed in aqueous McIlvaine’s buffers, except for the Yellow series where 30 % acetonitrile was added.

The BODIPY scaffold was chosen for its good photophysical properties and easily tunable emission wavelength. Red (**Red BODIpH**, λ_em_ = 597 nm) and far-red (**fr BODIpH**, λ_em_ = 665 nm) emitting derivatives were obtained by extending the electronic conjugation on position 3 and 5 with sulfonated styryl moieties (**Figure 2A&B**). BODIPYs display robust fluorescent properties but are intrinsically highly lipophilic and different strategies have been implemented to increase their solubility in aqueous media.^[24–27]^ Here the introduction of sulfonated moieties not only shifts the emission wavelength but also make them hydrosoluble to avoid aggregation and enable pH sensing at the interface of the particles with the aqueous environment.^[28]^

Absorption measurements conducted within the concentration range of 1–20 µM revealed that both the **Yellow BODIpH** and **Red BODIpH** series adhere to the Lambert–Beer law up to approximately 15 µM, whereas the **fr BODIpH** series exhibit no detectable solubility limit (Figure S1). Despite their apparent solubility, **Yellow BODIpH 1** and **2** display markedly reduced fluorescence in aqueous media compared to their sulfonated red and far-red analogues, indicative of aggregation-induced quenching. The use of aqueous buffers supplemented with 30 % acetonitrile allowed recovering the fluorescence and performing the photophysical characterization (Figure S2). The moderate solubility observed for the Yellow series, which lack solubilizing substituents apart from a short PEG chain, aligns with the intrinsic lipophilicity commonly associated with BODIPY derivatives. The red and far red BODIpH series could be characterized in fully aqueous media.

The extinction coefficients are high (approx. 60 000 cm^-1^mol^-1^L) and similar for the **Yellow** and **fr BODIpH** series but are surprisingly decreased for the **Red BODIpH** series (approx. 30 000 cm^-1^mol^-1^L, **Table 1**). The extension of conjugation leads to a decrease of quantum yield that can be due to the increased flexibility of the styryl branch that can favor non radiative decay (ϕ_F_ around 80 % for the yellow series, 50 % for the red series, 15 % for the fr series, **Table 1**). Overall, the brightnesses remain very satisfactory, especially for fully hydrosoluble BODIPYs with far red emissions.^[29,30]^

**Table 1.** Photophysical properties of the BODipH probes. Maximum absorption (λ_abs_) and emission (λ_em_) wavelength, molar extinction coefficient (ε), fluorescent quantum yield (ϕ_F_).

| | pH | $\lambda_{abs}$<br>(nm) | $\epsilon$ (cm<sup>-1</sup><br>mol<sup>-1</sup>L) | $\lambda_{em}$<br>(nm) | $\phi_F^a$ | $pK_a$ | pH sensing<br>range | Full<br>dynamic<br>range | Dynamic range<br>(pH4.5/pH7.5) |
| --- | --- | --- | --- | --- | --- | --- | --- | --- | --- |
| Yellow<br>BODipH 1 | 6.1 | 524 | 74000 | 538 | 0.74 | 7.7 ±<br>0.01 | 6 – 10 | 170 | NA |
|  | 10.8 | 520 | 76000 | 538 | 0.06 |  |  |  |  |
| Yellow<br>BODipH 2 | 3.8 | 527 | 56000 | 541 | 0.85 | 5.8 ±<br>0.08 | 5 – 6.5 | 90 | 64 |
|  | 7.4 | 522 | 56000 | 540 | 0.05 |  |  |  |  |
| Red<br>BODipH 1 | 6.1 | 577 | 36000 | 594 | 0.55 | 7.9 ±<br>0.05 | 6.8 – 9.6 | 80 | NA |
|  | 10.8 | 570 | 32000 | 594 | 0.04 |  |  |  |  |
| Red<br>BODipH 2 | 3.8 | 578 | 30000 | 597 | 0.40 | 5.9 ±<br>0.06 | 4.5 – 7.4 | 102 | 17 |
|  | 7.4 | 574 | 33000 | 597 | 0.02 |  |  |  |  |
| Fr BODipH<br>1 | 6.1 | 639 | 78000 | 662 | 0.15 | 8.1 ±<br>0.05 | 6.8 – 9.7 | 70 | NA |
|  | 10.8 | 632 | 70000 | 662 | 0.00<br>2 |  |  |  |  |
| Fr BODipH<br>2 | 3.8 | 642 | 70000 | 665 | 0.15 | 6.3 ±<br>0.08 | 4.8 – 7.8 | 150 | 10 |
|  | 7.4 | 637 | 75000 | 665 | 0.00<br>8 |  |  |  |  |

<sup>a</sup> Fluorescence quantum yields were measured in the most acidic pH indicated in the table for each probe.

The photophysical and pH-sensing properties summarized in **Figure 2, Table 1** and **Figure S3 s**how that all 6 probes display a pH-sensitive emission with excellent dynamic ranges (70 to 200-fold fluorescence increase in fluorescence intensity between high and low pH conditions). The p*K*_*a*_ is mostly governed by the structure of the phenol group and regardless of the emission wavelength of the fluorophore part similar p*K*_*a*_ are obtained for each series (1 and 2). A slight p*K*_*a*_ increase of about 0.5 pH unit is nonetheless observed upon increase of the fluorophore conjugation for the two series (Table 1). While these results provide insights into the design of tunable fluorescent pH probes, only the probes from the second series with p*K*_*a*_ values around 6 are useful to sense pH during endocytic processes. In addition to the full dynamic range between the most acidic and basic values, the dynamic range between pH 4.5 and pH 7.5 was also calculated for series 2 as it corresponds to the pH range involved during phagocytosis. **Red** and **fr BODIpH 2** display gradual pH transitions that slightly limit their dynamic range compared to **Yellow BODIpH 2** but leads to wider pH sensing range between 4.5 and 7.4 that is perfectly suited to monitor pH during endocytic processes such as phagocytosis. Most pH probes dedicated to endocytic processes are limited to narrower sensing ranges of two pH units around p*K*_*a*_.^[24,31,32]^

### Formulation of ratiometric pH-sensitive and targeted fluorescent microparticles

We previously reported the use of micrometer-sized oil-in-water emulsion droplets as biomimetic particles to study phagocytosis. These droplets can be readily functionalized at their surface with phagocytic ligands such as immunoglobulin G (IgG) or mannose and are efficiently recognized and internalized by macrophages in a receptor-selective manner.^[15,16,18]^ Building on these results, we adapted this platform to incorporate pH-sensitive probes for monitoring phagosomal maturation.

To formulate microparticle sensors, **fr BODIpH 2** was chosen for its sensing properties and far red emission that facilitates multiplexed imaging experiments. Thanks to its azide handle, it was conjugated to a DSPE lipid bearing a dibenzocyclooctyne (DBCO) group via copper-free azide–alkyne cycloaddition, yielding the amphiphilic compound **LipH** (**Figure 3A** and **Scheme S3**). **LipH** includes a long polyethylene glycol PEG_100_ spacer designed to minimize electrostatic interactions between the negatively charged particle surface and the pH-sensitive fluorophore. In presence of the droplets, the amphiphilic lipid spontaneously localizes at the oil–water interface, forming a stable pH-responsive fluorescent surface (**Figure 3B**,**C**).

**Figure 3.**
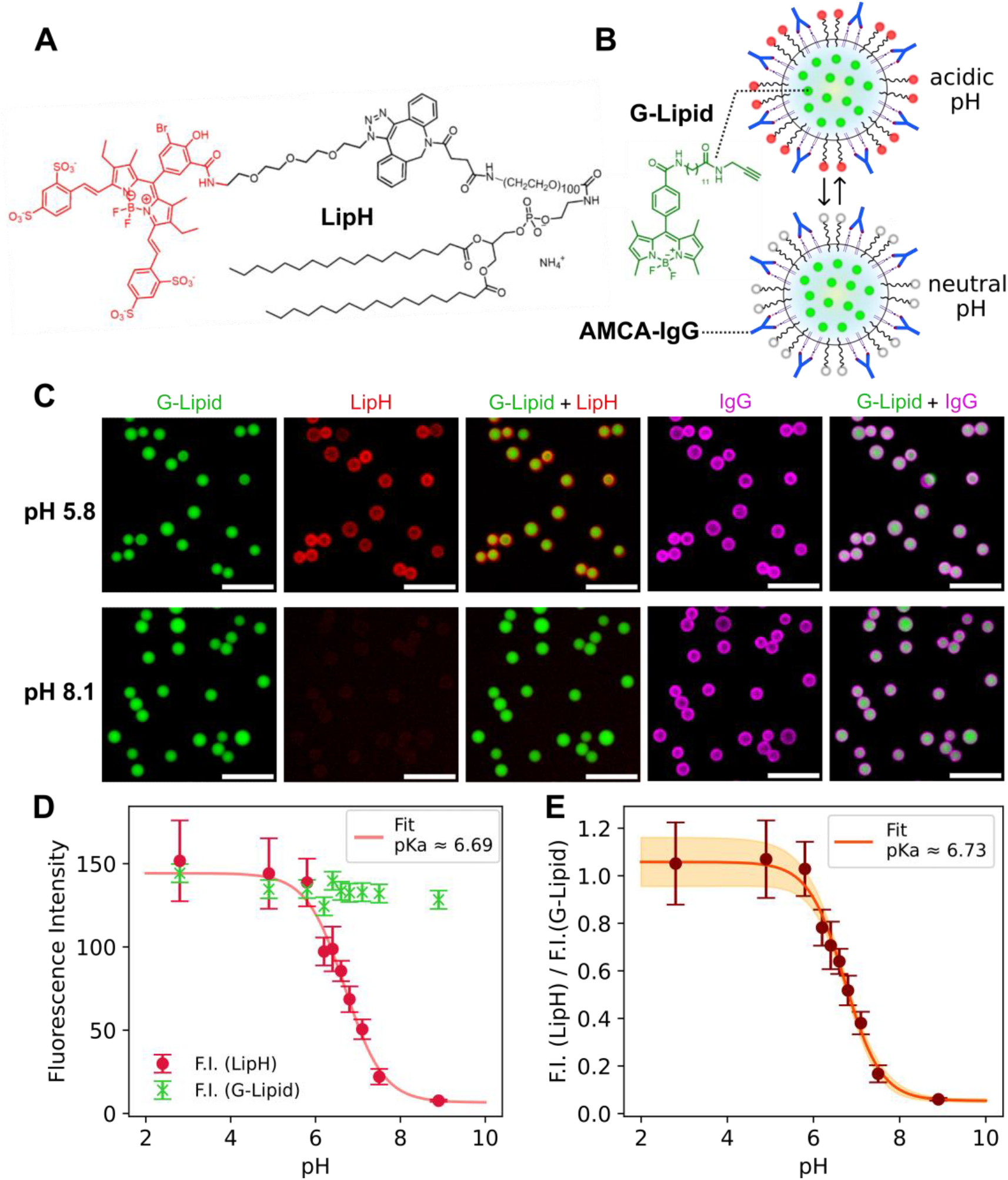
Design and characterization of ratiometric pH-sensitive droplets. (A) Chemical structure of the fluorescent pH-sensitive lipid **LipH**. (B) Schematic of the targeted ratiometric droplet design, with LipH and IgG positioned at the surface and a pH-insensitive reference dye (**G-lipid**) incorporated in the droplet core. (C) Confocal images of droplets co-functionalized with G-Lipid ( λ_ex_ = 488 nm / λ_em_ = 495–555 nm), IgG-AMCA (λ_ex_ = 405 nm / λ_em_ = 417–477 nm), and LipH ( λ_ex_ = 640 nm / λ_em_ = 645–725 nm) recorded at two different pH values. Scale bar: 30 µm. (D) Fluorescence intensities in the green (**G-lipid**) and red (**LipH**) channels of droplets functionalized with **LipH** and **G-Lipid** depending on pH. The red curve represents a sigmoidal fit of the red channel intensity, yielding a pKa of 6.69. (E) Ratio of the **LipH** fluorescence to the **G-Lipid** fluorescence as a function of pH. The red curve represents a sigmoidal fit, yielding a pKa of 6.73.

To achieve quantitative ratiometric measurements, we introduced a pH-insensitive reference dye into the droplet core. For this purpose, we used **G-lipid**—an intermediate in the synthesis of previously reported green-emitting fluorescent mannolipids—which, being non-amphiphilic, preferentially partitions into the oil phase of the droplets (**Figure 3B,C**).^[18]^ This internal standard, whose fluorescence remains constant under varying environmental conditions, allows normalization of the intensity changes from the surface-bound pH sensors.

To specifically engage Fcγ receptors on macrophages and trigger receptor-mediated phagocytosis, we co-functionalized 8 µm droplets with **LipH, G-lipid**, and AMCA-labeled IgG (λ_abs_ = 344 nm, λ_em_ = 440 nm) following a previously established protocol (Figure S5, **Figure 3B**).^[17]^

We calibrated the response of the functionalized droplets in solutions of cell culture medium (RPMI) whose pH were adjusted over a 2–9 range. The fluorescence was measured by epifluorescence microscopy in the far-red (**LipH**) and green (**G-lipid**) channels. The functionalized droplets exhibited excellent pH-sensing behavior closely matching that of the parent probe **fr BODIpH 2**, with a dynamic range showing nearly a 50-fold increase in fluorescence. The apparent p*K*_*a*_ was however slightly higher on the droplet surface (6.7) than in solution (6.3) leading to a sensing range between pH 5.0 and 8.0 (**Figure 3C**,**D**). In contrast, the fluorescence of **G-lipid** remained essentially constant across the entire pH range, providing a reliable internal reference for constructing calibration curves and enabling quantitative pH measurements (**Figure 3E**).

### Application to quantitative measurements of pH during phagocytosis

We next assessed the ability of these pH biosensors to report phagosomal acidification. When functionalized droplets were presented to RAW 264.7 macrophages, internalized ones appeared markedly brighter than those remaining extracellular (**Figure 4A**). Quantitative analysis of individual uptake events showed that **LipH** fluorescence increased strongly - by approximately 500–700% - immediately following closure of the actin cup, revealing that acidification begins as soon as the droplet is fully enclosed (**Figure 4B, Movie S1**). In some cases, transient fluctuations (“flickering”) precede stabilization (**Figure 4B, Figure S6A**).

**Figure 4.**
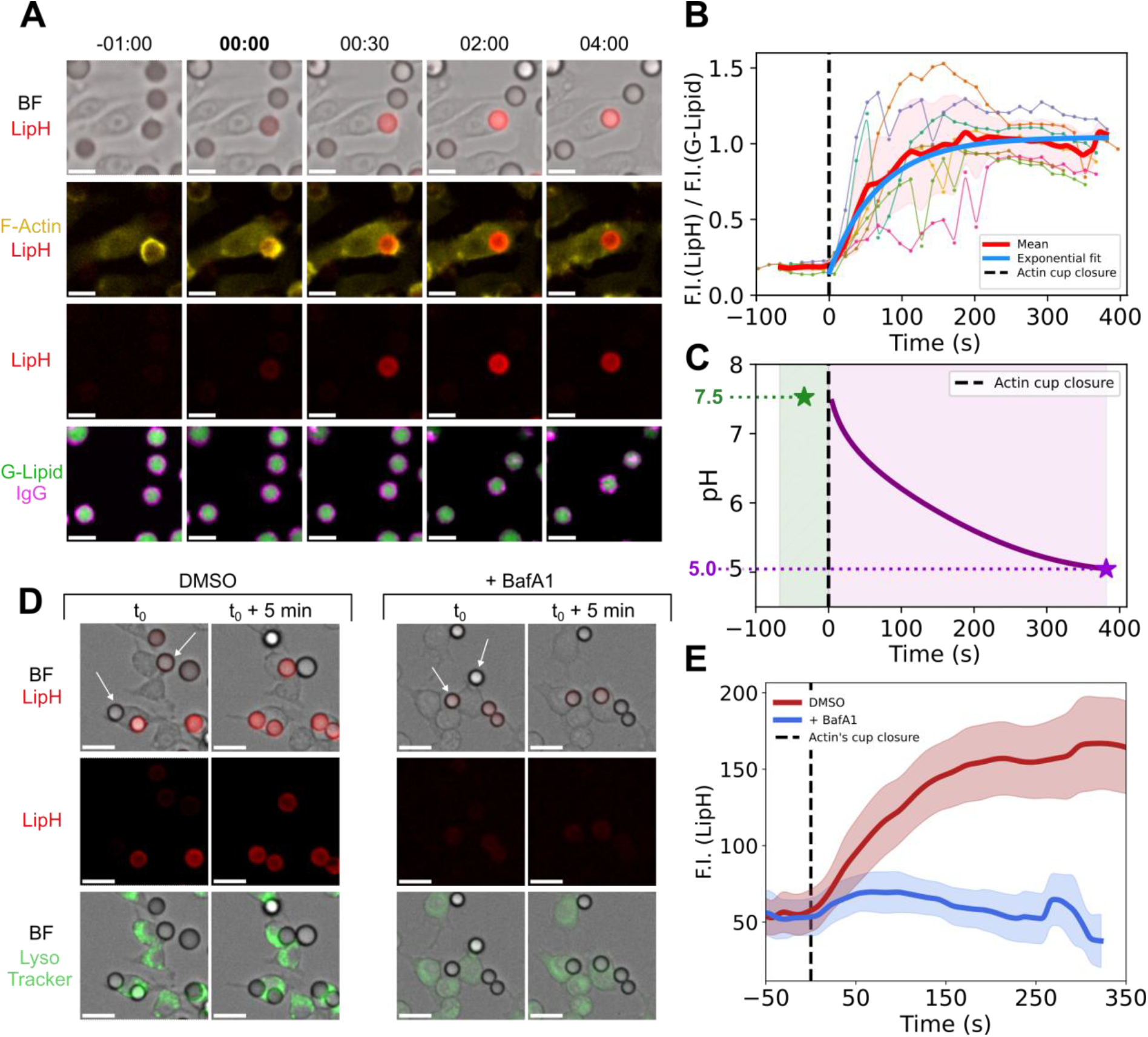
Quantitative phagosomal pH monitoring during the maturation stage. (A) Confocal time-lapse images of a lipid droplet co-functionalized with LipH (λ_ex_ = 640 nm / λ_em_ = 645–725 nm), G-Lipid (λ_ex_ = 488 nm / λ_em_ = 495–555 nm), and IgG-AMCA (λ_ex_ = 405 nm / λ_em_ = 417–477 nm) during internalization by a RAW264.7 macrophage. The F-actin channel (λ_ex_ = 561 nm / λ_em_ = 571–643 nm) marks the closure of the actin cup, indicating the moment of complete engulfment. Scale bar: 10 µm. (B) Time evolution of the LipH/G-Lipid fluorescence ratio for six individual droplets during internalization. Traces are aligned to the time of actin cup closure (t = 0); the mean trajectory is shown in red. In blue: exponential fit of the mean ratio after internalization. (C) Time evolution of the pH reported by the probe during internalization. In light green zone: extracellular pH estimated from the mean ratio at t<0 s. In light pink zone: intracellular pH curve durin maturation derived from the exponential fit of the ratio. (D) Confocal images of a lipid droplet functionalized with LipH and IgG in contact with RAW264.7 macrophages. Images were acquired at an initial time t_0_ and at t_0_ + 5 min. Droplets being internalized are indicated with a white arrow. On the left: 1 µL of DMSO was added with the cells for the control. On the right: cells were treated with BafA1 (200 nM) for 4 hours to prevent acidification. For both cases: lysosomes were stained with LysoTracker green (100 nM). Scale bar = 15 µm. (E) Time evolution of the mean fluorescence intensity for several droplets during internalization. In blue : the mean profile for macrophages treated with BafA1 (200 nM). In red : the mean profile for untreated macrophages. Traces are aligned to the time of actin cup closure (t = 0).

In contrast, G-lipid fluorescence decreased slightly over time due to photobleaching. This effect was corrected using non-internalized reference droplets, as detailed in the Supplementary Information and shown in **Figure S7**. After correction, **G-lipid** intensity could be considered constant and independent of pH variations (**Figure S6B)** . Consequently, the ratio of **LipH** to **G-lipid** fluorescence provides a reliable output to quantitatively measure phagosomal pH over time.

The averaged plot of the LipH/G-lipid fluorescence ratio was fitted with a single-exponential function, yielding a time constant τ = 69 s that reflects the kinetics of acidification (**Figure 4B**). Using the calibration curve (Figure 3E), the fitting function was converted to absolute pH values. The phagosomal pH decreased from an initial value of approximately 7.5—consistent with extracellular conditions—to about 5.0 within a little more than six minutes following actin cup closure (**Figure 4C**)

To confirm that this increase was specifically due to proton accumulation, the main mediator of phagosomal acidification, V-ATPase, was inhibited with Bafilomycin A1 (BafA1).^[3]^ Lysosomal fluorescence drastically decreased upon BafA1 treatment, reflecting the loss of acidification in phagosomal and lysosomal compartments (**Figure 4D, Figure S8**). This decrease confirms that the inhibitor effectively blocked V-ATPase activity, preventing LysoTracker accumulation.^[33]^ In this case, the droplet fluorescence in the red channel exhibits negligible enhancement after internalization relative to samples without the inhibitor, which supports the specificity of the probe (**Figure 4E, Movie S2, Movie S3**).

The kinetics of phagosomal acidification observed in our experiments are consistent with several reports in the literature, although discrepancies remain regarding the precise timing of pH transitions. In our experiments, IgG-functionalized droplets exhibited a rapid and immediate decrease in intraphagosomal pH, from approximately 7.5 before internalization to ∼5 within less than 10 minutes after actin cup closure. It should be noted that the dynamic range of our sensor is limited to pH values above 5 and does not allow monitoring the very final stage of phagosome maturation down to pH 4.5. Similar fast kinetics have been reported previously, where phagosomes were shown to acidify immediately following ingestion and to reach a minimal pH of around 5 within 10 – 15 minutes. ^[7,8]^

However, a recent study describes a three step trajectory: an initial neutral “stand-by” phase near-neutral pH, a rapid acidification period over a few minutes, and a plateau at pH 4.5-5.0.^[9]^ Our results therefore align with the faster end of this range of kinetics while underscoring the context-dependence of phagosomal maturation.

Our novel pH biosensors provide such control by allowing precise tuning of the particule size and functionalization to engage specific receptors and then follow phagosome maturation with high dynamic pH probes.

## CONCLUSION

In this study, we developed a ratiometric pH-sensing system based on BODIPY-derived probes incorporated into biomimetic lipid droplets. The hydrosoluble BODIpH fluorophores display tunable emission wavelengths and pKa values, enabling selective monitoring of acidification events in the physiologically relevant range. Incorporation of a pH-agnostic G-lipid reference allowed quantitative ratiometric readouts. When coupled to IgG, these droplets were efficiently internalized by macrophages via Fcγ receptors and reported rapid phagosomal acidification with high temporal resolution. The fluorescence increase of 500– 700% and stabilization of pH within approximately 6 minutes after actin cup closure indicate that the early stages of phagosome maturation may proceed faster than previously described. The final phagosomal pH of ∼5 is consistent with the establishment of an acidic environment for enzymatic activation, although values below pH 5 cannot be probed due to the intrinsic pKa of the sensor.

The development of fluorophores with lower p*K*_*a*_ values would extend the sensing range of the droplets and allow measurement of later stages of phagosomal maturation where pH decreases below 5. Thanks to the tunable structure of the BODIpH it should also be possible to combine several pH probes with distinct p*K*_*a*_ and emission wavelength to cover the whole range of phagosomal pH from pH 7.4 to 4.5 with high sensitivity.

Finally, the modularity of the oil-in-water droplets system should enable the incorporation of different targeting ligands, making it possible to investigate how distinct receptor pathways regulate acidification kinetics and steady-state pH but also other fluorescent probes for enzymatic activity or reactive oxygen species to investigate other parameters of phagosome maturation.

## ACKNOWLEDGMENTS

This work was supported by the Agence Nationale de la Recherche (ANR-20-CE13-0017) and benefited from the technical resources of the joint service unit CNRS UAR 3750 at the Institut Pierre-Gilles de Gennes. We thank Zoher Gueroui and Emma Pasquier (CPCV, Département de Chimie, ENS) for providing Bafilomycin A1 and P. Jurdic and K. Pernelle (IGFL, Lyon) for the Lifeact-mCherry RAW 264.7 macrophage cell line. We are also grateful to Guerbet (Villepinte, France) for supplying Lipiodol.

## DATA AVAILABILITY STATEMENT

The data that support the findings of this study are available from the corresponding author upon reasonable request.

## AUTHOR CONTRIBUTION

Sophie Michelis: Investigation, Synthesis, Data curation, Formal analysis, Visualization, Writing – original draft (figure and data preparation).

Héloïse Uhl: Investigation, Methodology (phagocytosis and imaging assays), Data curation, Formal analysis, Visualization, Writing – original draft.

Florence Niedergang: Funding acquisition, Supervision, Writing – review & editing.

Jacques Fattaccioli: Conceptualization, Supervision, Formal analysis, Visualization, Writing – original draft, Writing – review & editing, Funding acquisition.

Blaise Dumat: Conceptualization, Supervision, Formal analysis, Visualization, Writing – original draft, Writing – review & editing.

Jean-Maurice Mallet: Conceptualization, Supervision, Formal analysis, Writing – review & editing, Funding acquisition.

