## SupplementaryInformation for "Targeted fluorescent lipid microparticles for quantitative measurement of phagosomal pH"

### I. Synthesis of the BodipH and LipH probes

The synthetic pathway starts from a 3-carboxy-4-hydroxybenzaldehyde precursor (Scheme S1). The acid function was coupled to an amino-PEG$_3$-azido **2** prior to BODIPY synthesis. Initial synthesis efforts to form the **Yellow BODIpH 1** yielded a modest 10% yield with dichloromethane as solvent (procedure 1, Scheme S1). We sought to optimize the procedure for the BODIPY synthesis and obtained good results with the incorporation of ethanol as a co-solvent inspired by a procedure from Radunz *et al.*[1] This minor alteration resulted in a remarkable enhancement of the synthetic yield up to 70 % for **Yellow BODIpH 1** (procedure 2, scheme S1).

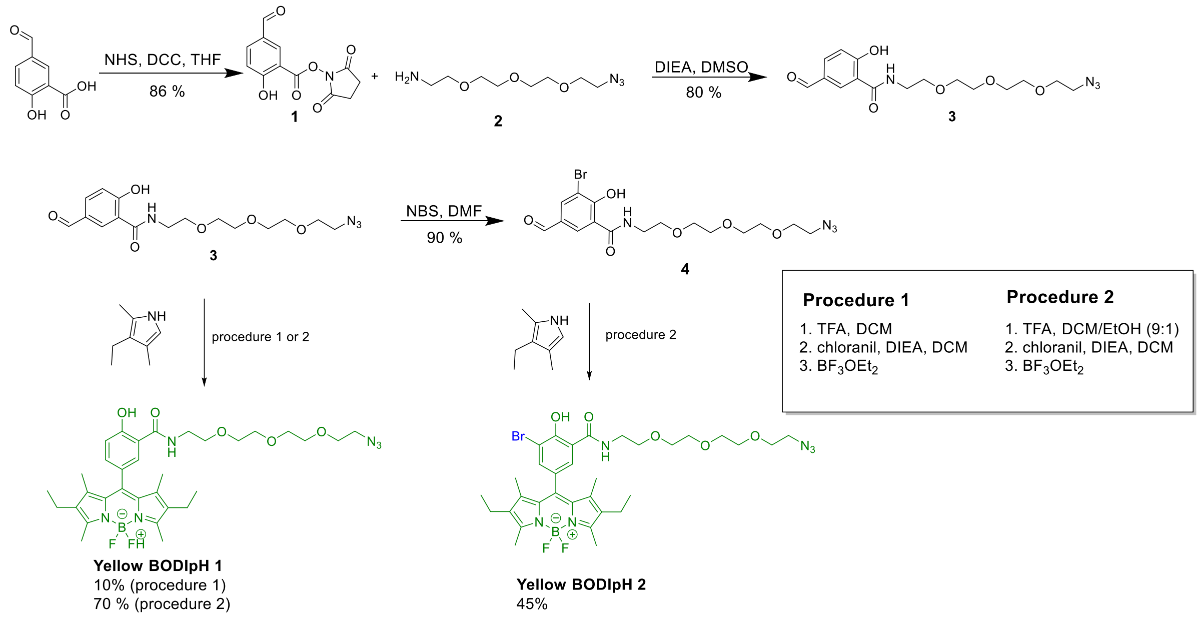

Scheme S1: Synthesis of yellow BODIpH.

To further decrease the p$K_a$ of our pH probes, we introduced a bromine atom in the other ortho position of the hydroxyl group. It is recognized that the incorporation of a bromine onto a phenolic structure leads to a reduction of its p$K_a$ by 1 to 2 units thanks to its attractive inductive effect.[2,3] Bromination was performed on the aldehyde prior to the BODIPY synthesis using N-bromosuccinimide (NBS) to obtain intermediate **4** with 90 % yield. **Yellow BODIpH 2** was then obtained in 45 % yield using the optimized procedure 2 (Scheme 1). It is worth noting that procedure 1 did not allow obtaining **Yellow BODIpH 2** at all, even in low yield.

To tune the emission wavelength and obtain hydrosoluble pH probes, we extended the conjugation on the methyl group in position 3 and 5 using Knoevenagel condensation with 4-formylbenzene-1,3-disulfonate (Scheme 2).

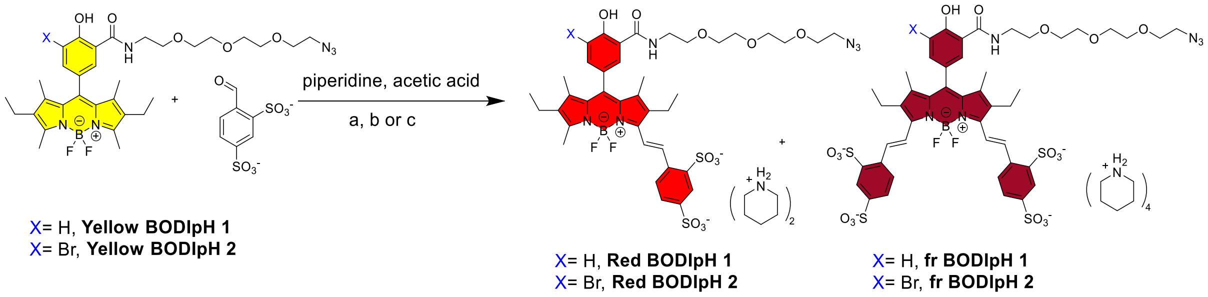

Scheme S2: Synthesis of **BODipH red 1**, **BODIpH red 2**, **BODIpH fr 1**, **BODIpH fr 2**. Conditions a, b, c from Table S1.

A procedure reported by Hoogendoorn *et al.*[4] with ethanol as a solvent and piperidine and acetic acid in large excess using microwave heating at 130°C afforded **Red BODIpH 1** and **fr BODIpH 1** starting from **Yellow BODIpH 1** but the yields were very low and we sought to optimize reaction conditions (Table S1).

Table S1: Overview of the reaction parameters for the coupling with 4-formylbenzene-1,3-disulfonate presented Scheme S2.

| | **Starting material** | **Solvent** | **Temperature** | **Time** | **Yield** |
|---|---|---|---|---|---|
| **a** | **Yellow BODipH 1** | Ethanol | 130 °C | 20 min | 1% **Red BODIpH 1**<br>5% **fr BODIpH 1** |
| **b** | **Yellow BODipH 1** | DMF | r.t. | 24h | 10% **Red BODIpH 1**<br>23% **fr BODIpH 1** |
| **b** | **Yellow BODipH 2** | DMF | r.t. and 25 °C | overnight | No product |
| **c** | **Yellow BODipH 2** | DMSO | 25 °C | 24h | 3% **Red BODIpH 2**<br>31% **fr BODIpH 2** |

The aldehyde is insoluble in methanol and acetonitrile even at ebullition temperature but seemed to be more soluble in dimethylformamide a solvent in which we managed to reach

33 % of conversion yielding the mono (10 % yield) and di-substituted (23 % yield) products with piperidinium counterions (Table 1). Due to the difficulty to stop the reaction after the mono-substitution, compounds from the Red BODIpH series are obtained as intermediary side-products of the bi-substitution with lower yields than for the fr BODIpH series. The polar and hydrosoluble products were purified by flash chromatography using a reverse phase C18 column. We performed the same reaction with **Yellow BODIpH 2**. Unfortunately, TLC monitoring showed that the reaction did not start even after 24 h using DMF as solvent, probably due to the lack of solubility of **Yellow BODIpH 2** in DMF. The latter was replaced by dimethylsulfoxide (DMSO), a solvent in which both reactants are soluble, and the conversion reached 34% at 25°C (Table 1, Scheme 2).

The pH-sensitive fluorescent lipid **LipH** was obtained by copper-free azide alkyne cycloaddition in DMF starting from fr BodipH 2 and DSPE-DBCO commercial lipids (Avanti research). The products were purified by size exclusion chromatography.

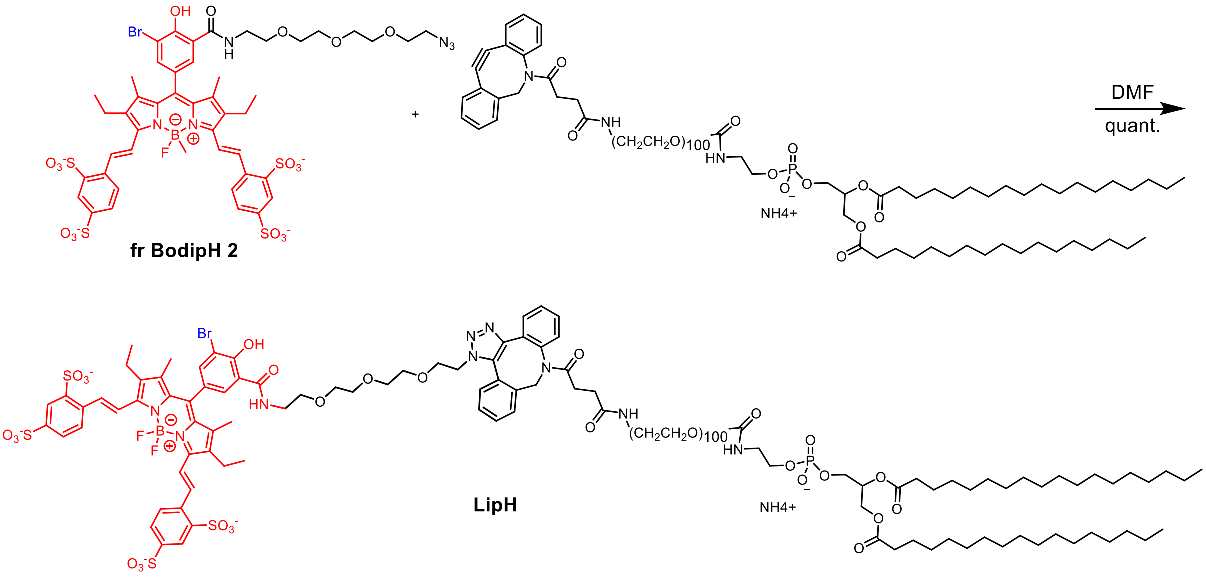

Scheme S3: Synthesis of **LipH**

#### II. Additional figures

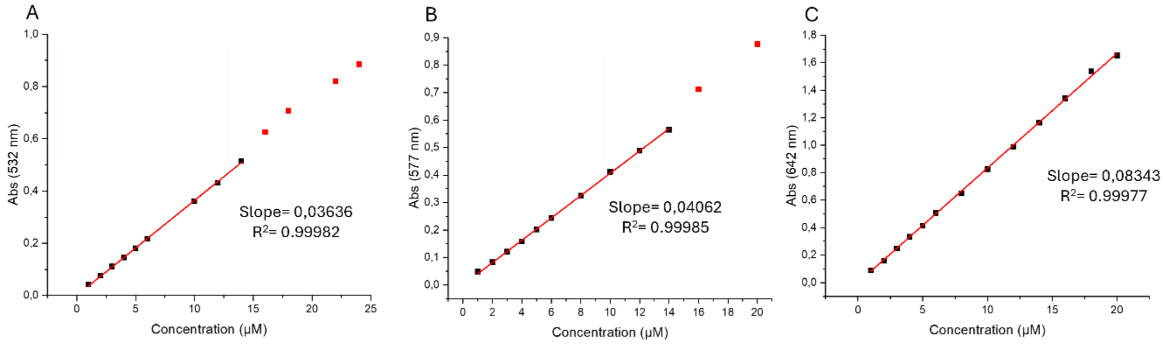

Figure S1: Absorbance of Yellow BODIpH 2 (A), Red BODIpH 2 (B), fr BODIpH 2 (C) with varying concentrations in water.

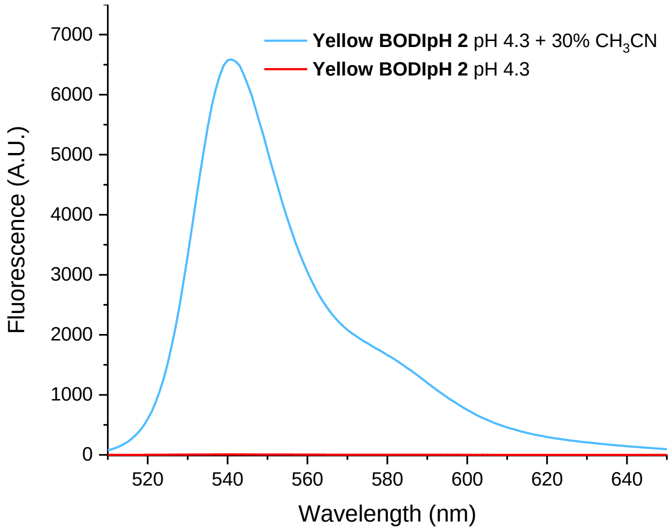

Figure S2. Fluorescence spectra of a 5 µM solution of Yellow BODIpH 2 in a pH 4.3 aqueous buffer with and without 30 % of acetonitrile.

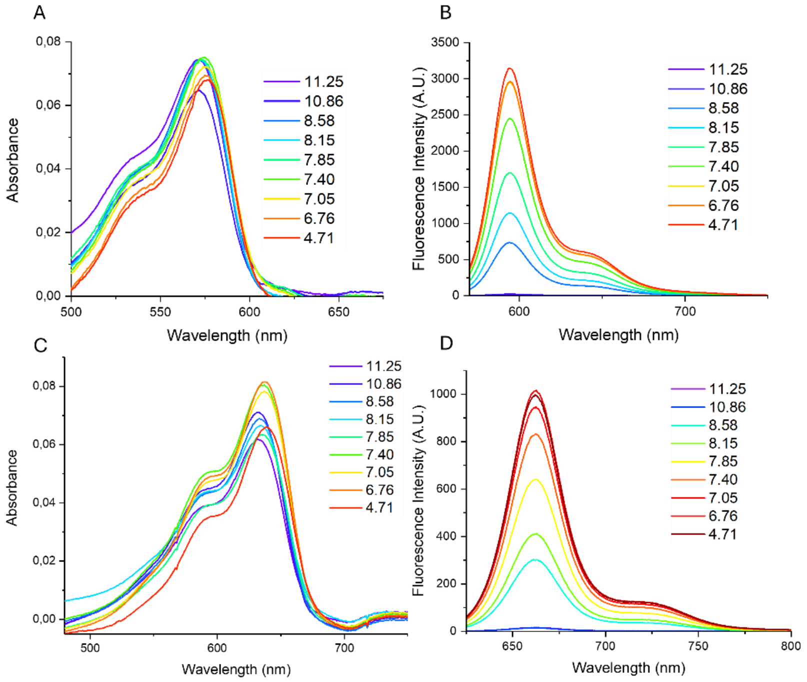

Figure S3 (A) Absorbance of **Red BODIpH 1** in Britton-Robinson buffers, (2 µM). (B) Fluorescence emission of **Red BODIpH 1** in Britton-Robinson buffers, (2 µM), $\lambda_{ex}$= 560 nm(C) Absorbance of **fr BODIpH 1** in Britton-Robinson buffer. (1 µM) (D) Fluorescence emission of **fr BODIpH 1** in Britton-Robinson buffers. $\lambda_{ex}$= 620 nm. (1 µM)

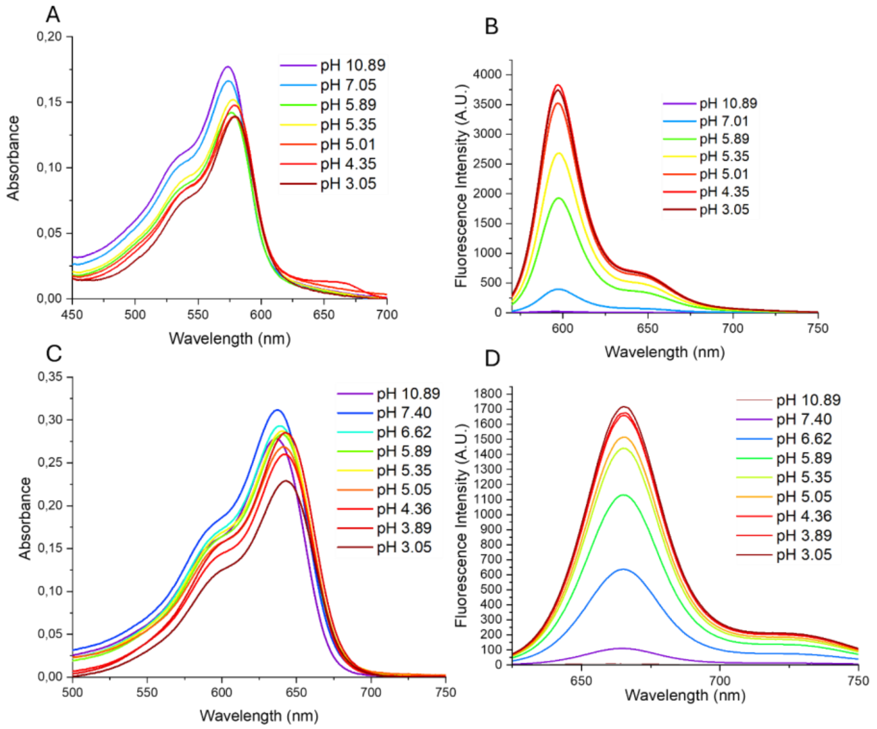

Figure S4: (A) Absorbance of **Red BODIpH 2** in Britton-Robinson buffer. (5 µM) (D) Fluorescence emission of **Red BODipH 2** in Britton-Robinson buffers. $\lambda_{ex}$ = 560 nm. (5 µM). (C) Absorbance of **fr BODIpH 2** in Britton-Robinson buffer. (4 µM) (D) Fluorescence emission of **fr BODIpH 2** in Britton-Robinson buffers. $\lambda_{ex}$ = 620 nm. (4 µM)

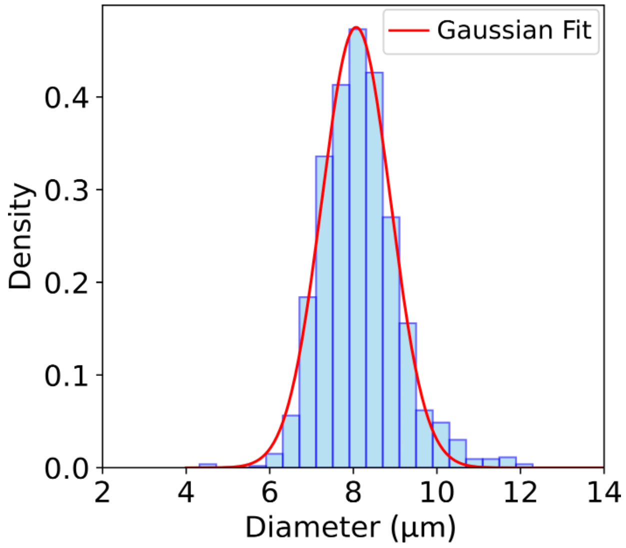

Figure S5. Size distribution of the lipiodol emulsion droplets having a mean diameter of 8.1 ± 0.8 µm. Average of 1337 droplets, determined with fluorescence microscopy.

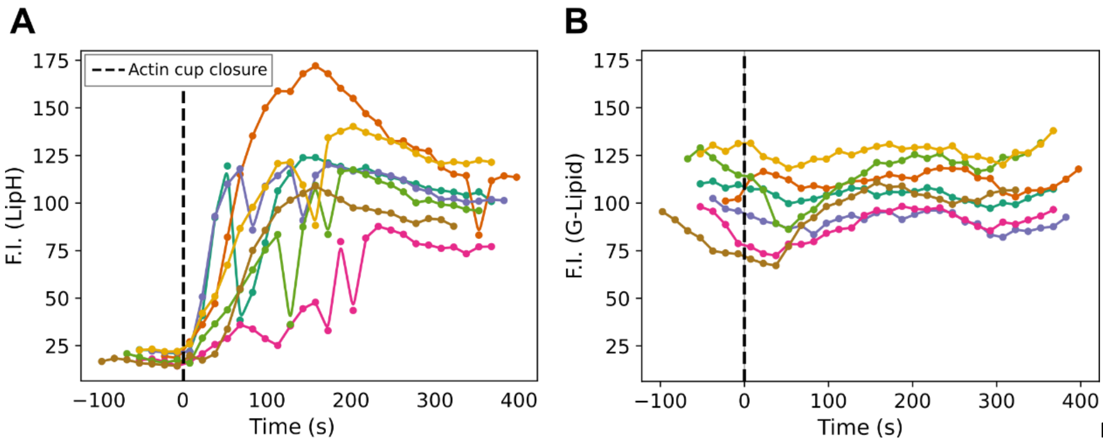

Figure S6: (A) Fluorescence in the LipH channel ( $\lambda_{ex}$ = 640 nm / $\lambda_{em}$ = 645–725 nm) of 7 different droplets during the process of internalization. (B) Fluorescence in the G-Lipid channel ($\lambda_{ex}$= 488 nm / $\lambda_{em}$= 495 - 555 nm) of 7 different droplets during the process of internalization.

#### Correction of the bleaching fluorescence

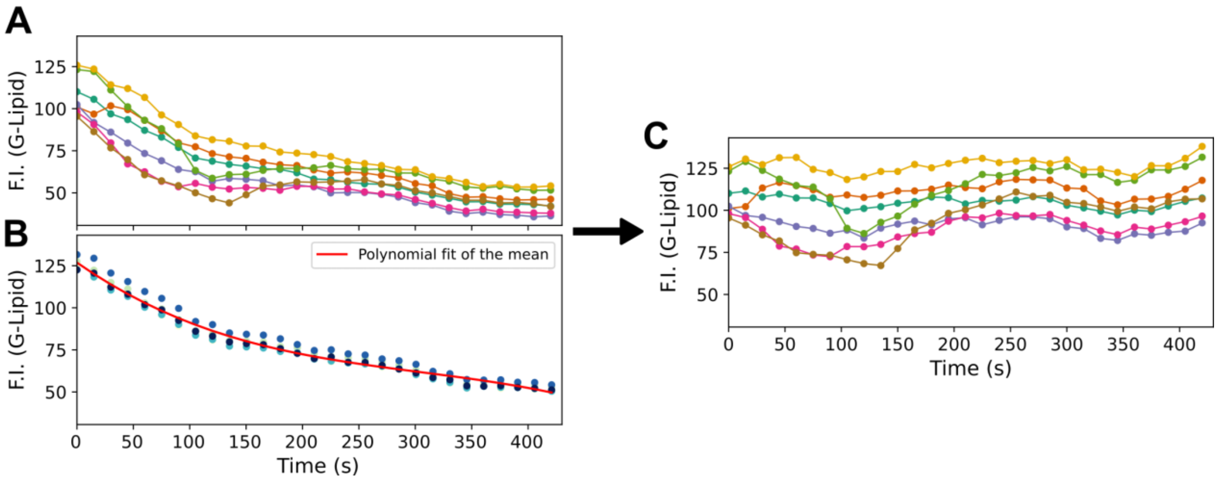

Figure S7: (A) Fluorescence in the G-Lipid channel ($\lambda_{ex}$= 488 nm / $\lambda_{em}$= 495 - 555 nm) of 7 different droplets during the process of internalization. (B) Fluorescence in the G-Lipid channel of 4 different droplets that were not internalized. The microscope settings and analysis routine used are the same as those used for (A). In red, polynomial fit of the mean intensity according time. (C ) Fluorescence in the G-Lipid channel of the same 7 droplets as (A) during the process of internalization after the bleaching correction.

To correct the G-Lipid's intensity fluorescence decrease due to bleaching, 4 droplets that remained not internalized were taken into account (**Figure S7,B**). The droplet's fluorescence intensity in the G-Lipid's channel ($\lambda_{ex}$= 488 nm / $\lambda_{em}$= 495 - 555 nm) was measured for each of them and the mean at each time was measured and fitted by a polynomial function *Bleach(t)* .

The intensity measured for the 6 droplets considered for the study was modified this way :

$$F.I._{GLip,corr}\,(t)=\frac{Bleach(0)}{Bleach(t)}\times F.I._{GLip}\,(t)$$

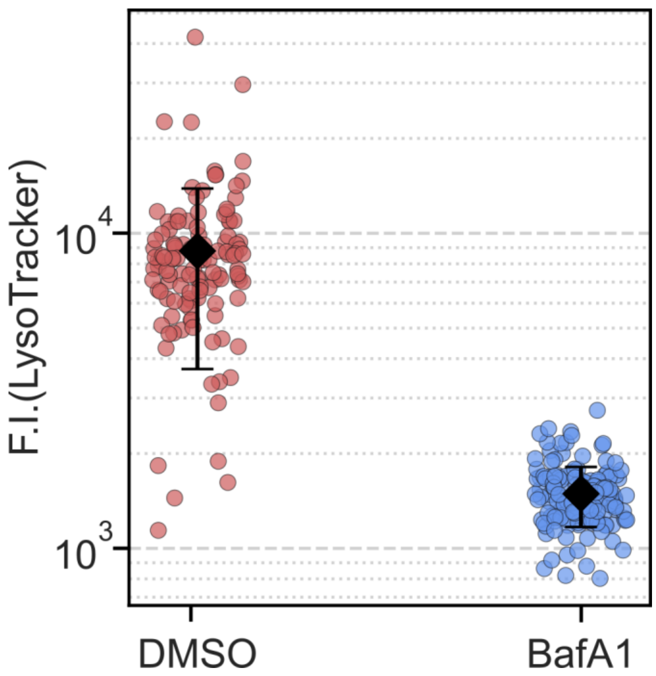

Figure S8 : Maximum fluorescence intensity in the green channel ($\lambda_{ex}$= 488 nm / $\lambda_{em}$= 495 - 555 nm, corresponding to the LysoTracker emission) measured in individual macrophages incubated with LysoTracker (100 nM) for 20 minutes. Two conditions were analyzed: control cells treated with 1 µL DMSO (in pink, n = 120) and cells treated with BafA1 (200 nM) for 4 hours (in blue, n = 153). The same imaging parameters were used for both cases. Each point represents a single cell, and the black diamond indicates the mean ± standard deviation.

#### III. Material and methods

All starting materials for the synthesis were purchased from Alfa Aesar, Sigma-Aldrich, or TCI Europe and used as received unless stated otherwise. NMR spectra were recorded on a Bruker Advance 300 MHz spectrometer. Mass spectra were obtained using a LTQ-Orbitrap XL from Thermo Scientific (Thermo Fisher Scientific, Courtaboeuf, France) and operated in positive or negative ionization mode, with a spray voltage at 3.6 kV. For reactions purposes, all solvents were used dried and purchased from Sigma-Aldrich.

**UV-Vis absorption and fluorescence spectroscopies**

Absorption spectra were recorded on a Cary 300 UV-Vis spectrophotometer with a 500 nm/s scan speed and a 1 nm step. Spectra were corrected with a baseline. The extinction coefficients were determined according to the Beer-Lambert law by measuring the absorption of solutions of increasing concentrations.

Emission spectra were recorded on a Jasco FP-8300 spectrofluorometer with a 600 nm/s scan speed and a 1 nm step. The sensitivity was set to medium, and the excitation and emission bandwidths were set to 2.5 nm.

Fluorescence quantum yields were measured by comparison to established reference Rhodamine 6G in ethanol ($\phi$ = 0.94).[5] For each dye, the absorbance and fluorescence of five solutions at increasing concentrations were measured. After plotting the curve of the area of fluorescence intensity against the absorbance I = f(A) at the excitation wavelength, the quantum yields $\phi_F$ were calculated using the following equation:

$$\phi_{sample} = \phi_{ref} \frac{grad_{sample}}{grad_{ref}} \times \frac{\eta^2_{sample}}{\eta^2_{ref}}$$

where grad is the slope of the I = f(A) curve and $\eta$ the refractive index of the solvent.

Fluorometric pH titrations curves were determined by measuring the fluorescence intensity of the probes at various pH by fluorescence spectroscopy. We used stock solution of the probes at 1 mM in DMSO. Britton-Robinson buffers of various pH were made by mixing various amount of 40 mM boric acid, 40 mM acetic acid and 40 mM phosphoric acid solutions and adjusting the solutions with hydrochloric acid or sodium hydroxyde.[6] For **BODIpH yellow 1** and **2** the measures were performed in 70% Britton-Robinson buffer and 30% acetonitrile whereas for **BODIpHs red and fr** the measures were performed in Britton-Robinson buffers only. The fluorescence of each probe was measured three times in each pH buffer. To determine the pKa, the integrated fluorescence intensity was plotted against pH and fitted using Hill1 equation in Origin 2018 software.

Solubility limits in water were determinde by measuring the absorbance of the probes at various concentrations in water. We used stock solution of the probes at 1 mM in DMSO. The absorbance at the maximum absorption wavelength for each solution was plotted against the concentration and fitted to a linear model.

**Lipid droplets formulation**

Droplets are fabricated with a Shirasu Porous Glass (SPG) membrane emulsification device. An hydrophilic-treated microporous membrane is immersed into the continuous phase of Poloxamer 188 (Sigma-Aldrich, Ref: P2443) at 15% w/w to remove the trapped air from the pores. The dispersed phase consisting in Lipiodol (Guerbet, (CAS no. 8002-46-8)) is inserted in a tank connected to a pressure valve. A pressure is applied to force the dispersed phase through the micrometer pores. At a critical pressure, droplets form, reach the continuous phase and are stabilized by surfactants. The resulting emulsion is continuously stirred during the process. Droplet's diameters depend on the pore size of the membrane. For the following experiment, a membrane of 2.1 µm was used to obtain droplets with well-controlled size diameter of 8 +/- 0.8 µm (Figure S5). The emulsion is stocked out of light at a temperature of 12°C in a Peltier-cooled cabinet.

**Droplets functionalization**

The volumes used for the surface functionalization are expressed as volume equivalent. One equivalent (eq.) corresponds to the volume of molecules necessary to cover the total area of a population of droplets with a compact monolayer of molecules.[7]

The functionalization of droplets was performed following this procedure :

10 µL of 8 µm-large droplets (i.e. 16 millions droplets) in Poloxamer 188 (Pluronic F68, Sigma-Aldrich), at 15% w/v are placed in a microtube (Axygen, 0.6ml Microtubes, "Maximum recovery", Ref : MCT-060-L-C) and are diluted with phosphate buffer (pH = 7.1, 20mM) supplemented with Tween20 at the CMC (0.007 % w/v) in a way to reach a volume of 200µL. The diluted emulsion was centrifuged at 2000 rpm for 30 sec. The continuous phase was removed and 195µL of PB/Tween20 CMC were added again. This washing step was repeated 3 times. After the last rinsing step, only 10µL of the emulsion volume remain. Then, 20 eq. of DSPE-PEG(2000)-Biotin (Avanti Lipids, Ref : 880129P-10mg) (i.e. 2.6µL for a concentration of 10mg/mL) and 2.5µL of G-Lipid (at 40 µM in DMSO) were added, followed by DMSO to reach a volume of 20µL of polar solvent. Then the mix was completed with PB/Tween20 CMC to have a total volume of 200µL. After 30 minutes rotation in the dark, droplets were washed 3 times with PB/Tween20 CMC. After the last rinsing step, all the supernatant is discarded. Then, 0.5 eq. of IgG anti-biotin (Jackson Immunoresearch, AMCA-conjugated IgG Fraction Monoclonal Mouse Anti-Biotin, Ref : 200-152-211) (i.e. 3.9µL at 0.85mg/mL) were added to the droplets and the volume is completed with 50µL PB/Tween20 CMC. The mixture was then placed on a rotary shaker for 30min in the dark and was washed again 3 times.

Finally, to functionalize the droplets with the pH-probe, 0.5eq. of the LipH (i.e. 3µL for a concentration of 0.4mg/mL) was added, followed by DMSO to reach a volume of 20µL of polar solvent, and completed with PB/Tween 20 CMC to have a total volume of 200µL.

After 30 minutes rotation in the dark, droplets were washed 3 times with PB/Tween 20 CMC and were ready to be put in contact with macrophages.

**Fluorometric pH titrations curves**

Fluorometric pH titrations curves were determined by measuring the mean fluorescence intensity of the droplets in medium of various pH. To have an environment similar to the one that is used for cell experiments, HCl or NaOH was added to the cell medium while the pH of the solution was controlled with a pH-meter.

The mean of fluorescence intensity in the LipH channel ( $\lambda_{ex}$ = 640 nm / $\lambda_{em}$ = 645–725 nm) and the ratio $F.I._{LipH}/F.I._{G\text{-}Lipid}$ were plotted according pH and were fitted using a sigmoïd equation :

$$f(pH)=\frac{L}{1+e^{-k(pH-pKa)}}+B$$

yielding the following parameters :

For $F.I._{LipH}(pH)$ : L = -137.69 ± 14.98 ; k = 2.21± 0.38 ; pKa = 6.69 ± 0.12 ; B = 144.27 ± 14.48

For $F.I._{LipH}/F.I._{G\text{-}Lipid}$ (pH) : L = -1.00 ± 0.11 ; k = 2.37 ± 0.42 ; pKa = 6.73 ± 0.11 ; B = 1.06 ± 0.11

**Fit of the mean ratio $F.I._{LipH}/F.I._{G\text{-}Lipid}$**

The averaged plot of the LipH/G-lipid fluorescence ratio was fitted with a single-exponential function : $R(t) = A(1-e^{-t/\tau})+B$

yielding the following parameters : A = 0.90 ± 0.01, τ = 69.1 ± 5.9 and B = 0.14 ± 0.01

**Microsopy**

To study internalization, about one droplet for two cells was used. After waiting 5 min to let the droplets reach the cells, the well was imaged in live at 37°C and under controlled-atmosphere (5% CO2).

Cells' experiments as well as the one that provides the titration curves were performed with a spinning-disk inverted microscope (Leica DMi8). Images were recorded with MetaMorph (Molecular Device). The steam mode with the following parameters were used :

4 Z-steps of 3 µm

For the pH-probe LipH : $\lambda_{ex}$= 640 nm / $\lambda_{em}$= 685±40nm

For the F-actin : $\lambda_{ex}$=561nm / $\lambda_{em}$= 607±36nm

For the G-Lipid :$\lambda_{ex}$=488nm / $\lambda_{em}$=525± 30nm

For the IgG : $\lambda_{ex}$=405nm / $\lambda_{em}$=525± 30nm

When timelapses were recorded, a step of 15 seconds was used.

#### Image analysis

Measurements of fluorescence intensity were performed on ImageJ (Fiji Distribution)[8] .

**Fluorescence intensity of LipH and G-Lipid** :

For the analysis of fluorescence channels, Z-stacks were combined using an average Z-projection, and background fluorescence was subtracted.

Since droplets were labeled with G-Lipid, their contours were clearly visible, allowing segmentation using a fixed threshold applied to the green channel. The resulting binary mask was used to measure the mean fluorescence intensity in each frame for the red channel (LipH signal) and the green channel (G-Lipid signal). The same binarization threshold was consistently applied to all droplets.

Actin cup closure was determined manually. The last frame in which the droplet appeared partially engulfed was defined as $t_{last,out}$ , and the first frame in which the droplet was fully internalized as $t_{first,in}$ . The time of cup closure was calculated as: $t_{closure}$ =( $t_{first,in}$-$t_{last,out}$)/2.

**Fluorescence intensity of LysoTracker** :

For LysoTracker quantification, Z-stacks in the green channel were combined using an average Z-projection, and background fluorescence was removed.

Lysosomal fluorescence was segmented by thresholding the green channel using a constant minimal threshold across all conditions. For each analyzed cell, the maximum fluorescence intensity was measured.

Data analysis and plots were performed with Python (v 3.9.13)

**Cell culture**

Phagocytic assays were performed in RAW 264.7 murine mouse macrophage cell line transfected with LifeAct mCherry. Cells are grown in 100 mm diameter Petri dishes (Falcon, Ref : 353003) at 37°C and in a humidified atmosphere containing 5% of CO2. Macrophages are plated in complete medium consisting of RPMI Glutamax™ (Gibco, Ref : 61870-010) supplemented with 10% Fetal Bovine Serum (Gibco, FBS One Shot™, Heat Inactivated, Ref : A3840402), 10mM HEPES (Gibco, Ref : 15630-056), 1mM sodium pyruvate (Gibco, Ref : 11360-070) , 50µM 2-Mercaptoethanol (Gibco, Ref : 31350-010)and 2mM L-glutamine (Gibco, Ref : 25030-024). The expression of Lifeact_mCherry fluorescent protein was maintained by intermittent addition of 4µg.mL$^{-1}$ of puromycin (Gibco, Ref : A11138-03).

24 hours before experiments, cells were detached with a scraper (Costar, Ref : 3008) and seeded in new medium on a 24 well glass plates (Cellvis, Ref : P24-1.5H-N) to reach a concentration of about 2,5.10$^{5}$ cells/well. Just before imaging, cell medium was replaced by the same medium free of FBS and red phenol (Gibco, Ref : 11835030).

**For Bafilomycin treatment** :

Cells were pretreated with Bafilomycin A1 (Sigma-Aldrich, Ref. 196000-10UG) at 200 nM for four hours. Twenty minutes prior to imaging, the culture medium was replaced with the same medium free of FBS and red phenol, supplemented with 100 nM LysoTracker green and 200 nM Bafilomycin A1. Cells were imaged directly in this treatment medium.

### IV. Synthetic procedures

##### Compound 1

Compound 1 was obtained according to literature with a 86% yield.[9]

$^{1}$**H NMR** (300 MHz, Chloroform-*d*) δ 10.04 (s, 1H), 9.91 (s, *J* = 0.6 Hz, 1H), 8.54 (d, *J* = 2.0 Hz, 1H), 8.13 (dd, *J* = 8.7, 2.1 Hz, 1H), 7.19 (d, *J* = 8.7 Hz, 1H), 2.95 (s, 4H).

##### Compound 3

To a solution of **1** (0.300 g, 1.14 mmol, 1 eq) in DMSO (6 mL) under argon was added 2-(2-(2-(2-azidoethoxy)ethoxy)ethoxy)ethan-1-amine (0.298 g, 1.37 mmol, 1.2 eq) and DIEA (0.490 mL, 2.85 mmol, 2.5 eq). The reaction mixture was stirred at r.t. for 4h, then diluted in EtOAc, washed with HCl 1M and extracted (x3). The organic layer was dried over $MgSO_4$, filtered and concentrated. The crude product was purified by flash column chromatography (Cyclohexane-EtOAc 60:40 to 40:60). The mixture was concentrated to obtain **3** as orange oil (0.253 g, 80 % yield).

$^{1}$**H NMR** (300 MHz, Chloroform-*d*) δ 13.34 (s, 1H), 9.88 (s, 1H), 8.13 (d, *J* = 2.0 Hz, 1H), 7.90 (dd, *J* = 8.6, 2.0 Hz, 1H), 7.45 (s, 1H), 7.09 (d, *J* = 8.6 Hz, 1H), 3.78 – 3.60 (m, 14H), 3.36 (m,2H).$^{13}$**C NMR** (75 MHz, Chloroform-*d*) δ 190.12, 169.40, 167.09, 135.77, 128.22, 127.82, 119.21, 114.71, 70.79, 70.74, 70.60, 70.34, 70.00, 69.31, 50.69, 39.77.**HRMS** (ESI) calculated for $C_{16}H_{21}N_4O_6$ 365.1467. Found 365.1464

##### Yellow BODIpH 1 procedure 2

To a solution of **3** (0.290 mg, 0.79 mmol, 1 eq) in dry DCM (40 mL) and ethanol (3 mL) under argon was added 3-ethyl,2,4-dimethyl-1H-pyrrole (0.195 mg, 1.58 mmol, 2 eq). 5 rapid vacuum-argon cycles were performed. TFA was added (9 µL). The reaction mixture was stirred overnight in the dark until completion. Chloranil (0.194 g, 0.79 mmol, 1 eq) was added and the reaction mixture was stirred for 1 hour. The mixture was concentrated under reduced pressure and the product was diluted in dry DCM (30 mL). DIEA (1.3 mL, 7.9 mmol, 10 eq) was added followed by $BF_3OEt_2$ (1.95 mL, 15.8 mmol, 20 eq). The reaction mixture was stirred 1 hour. The reaction mixture was washed with water (x2), extracted with DCM (x2). The organic layer was dried over $MgSO_4$, filtered and concentrated. The crude product was purified by a first short flash column chromatography (90:10 DCM-Cyclohexane to DCM) and then Cyclohexane-EtOAc 90:10 to Cyclohexane-EtOAc 60:40. The mixture was concentrated to obtain **Yellow BODIpH 1** as red solid (0.380 g, 70% yield).

$^{1}$**H NMR** (300 MHz, Chloroform-*d*) δ 12.79 (s, 1H), 7.41 (d, *J* = 2.0 Hz, 1H), 7.28 (dd, *J* = 8.5, 2.0 Hz, 1H), 7.22-7.18 (m, 1H), 7.11 (d, *J* = 8.5 Hz, 1H), 3.65-3.50 (m, 8H), 3.46-3.32 (m, 6H), 3.24 (t, *J* = 5.0 Hz, 2H), 2.52 (s, 6H), 2.31 (q, *J* = 7.6 Hz, 4H), 1.38 (s, 6H), 0.99 (t, *J* = 7.5 Hz, 6H). $^{13}$**C NMR** (75 MHz, Chloroform-*d*) δ 169.57, 162.25, 154.03, 138.99, 138.29, 133.92, 132.99, 131.13, 125.69, 119.41, 114.86, 70.68, 70.59, 70.38, 70.14, 69.72, 69.10, 50.61, 39.66, 26.92, 17.05, 14.65, 12.51, 12.22. **HRMS (ESI) m/z** [M+H]+: calculated for $C_{32}H_{42}BF_2N_6O_5$ 639.3283 Found 639.3288

##### Compound 4

A solution of **3** (O.240 g, 0.655 mmol, 1 eq) in DMF under argon was stirred in an ice bath. NBS (0.117 g, 0.66 mmol, 1.01 eq) was added slowly. The mixture was heated to 70 degrees for 3 hours. The reaction mixture was concentrated under reduced pressure, diluted in DCM and washed with thiosulfate. It was extracted with DCM, dried over $MgSO_4$, filtered and concentrated. The crude product was purified by flash chromatography with 0.1% TFA, Cyclohexane-EtOAc 50:50 to 30:70. The mixture was concentrated to obtain **4** as light brown oil (0.259 g, 90 % yield).

**$^1$H NMR** (300 MHz, Methanol-*d*$_4$) δ 9.70 (s, 1H), 8.19 (d, *J* = 2.0 Hz, 1H), 8.08 (d, *J* = 1.9 Hz, 1H), 3.58 (m, 14H), 3.27 (t, *J* = 4.9 Hz, 2H). **$^{13}$C NMR** (75 MHz, Methanol-*d*$_4$) δ 189.56, 179.35, 169.06, 137.37, 129,17, 127.82, 115.83, 113,36, 77.63, 77.20, 76.77, 70.50, 70.42, 70.40, 69.99, 69.92, 69.36, 50.55, 49.73, 49.44, 49.16, 48.87, 48.59, 48.30, 48.02, 39.72. **HRMS (ESI) m/z**: [M+Na]+ calculated for $C_{16}H_{21}BrN_4O_6Na$ 467.0537. Found 467.0534

**Yellow BODIpH 2**

To a solution of **4** (0.247 mg, 0.55 mmol, 1 eq) ) in dry DCM (35 mL) and ethanol (2.5 mL) under argon was added 3-ethyl, 2,4-dimethyl-1H-pyrrole (0.195 mg, 1.58 mmol, 2 eq). 5 rapid vacuum-argon cycles were performed. TFA was added (6 µL). The reaction mixture was stirred overnight in the dark until completion. Chloranil (0.135 g, 0.55 mmol, 1 eq) was added and the reaction mixture was stirred for 1 hour. The mixture was concentrated under reduced pressure and the product was diluted in dry DCM (30 mL). DIEA (0.950 mL, 5.5 mmol, 10 eq) was added followed by $BF_3OEt_2$ (1.36 mL, 11 mmol, 20 eq). The reaction mixture was stirred 1 hour. The reaction mixture was washed with water (x2), extracted with DCM (x2). The organic layer was dried over $MgSO_4$, filtered and concentrated. The crude product was purified by flash column chromatography (Cyclohexane-EtOAc 90:10 to Cyclohexane-EtOAc 70:30. The mixture was concentrated to obtain **Yellow BODIpH 2** as red solid (0.166 g, 45% yield).

**$^1$H NMR** (300 MHz, Chloroform-*d*) δ 13.77 (s, 1H), 7.59 (d, *J* = 1.9 Hz, 1H), 7.57-7.52 (m, 1H), 7.49 (d, *J* = 2.0 Hz, 1H), 3.73 – 3.53 (m, 8H), 3.53 – 3.44 (m, 2H), 3.45 – 3.35 (m, 4H), 3.28 – 3.18 (m, 2H), 2.51 (s, 6H), 2.31 (q, *J* = 7.5 Hz, 4H), 1.42 (d, *J* = 1.4 Hz, 6H), 0.99 (t, *J* = 7.5 Hz, 6H). **$^{13}$C NMR** (75 MHz, Chloroform-*d*) δ 169.09, 159.01, 154.44, 138.12, 137.24, 136.85, 133.22, 130.95, 126.38, 125.18, 115.75, 113.01, 77.47, 77.04, 76.62, 70.61, 70.49, 70.32, 70.05, 69.64, 69.04, 50.60, 39.94, 26.92, 17.05, 14.66, 12.55, 12.48. **HRMS (ESI) m/z:** [M+Na]+ Calculated for $C_{32}H_{42}BBrF_2N_6O_5Na$ 741.2353. Found 741.235

**Red and fr BODIpH 1**

To a solution of **Yellow BODIpH 1** (0.035 g, 0.055 mmol, 1 eq) in DMF (1 mL) under argon were 4-formylbenzene-1,3-disulfonate (0.102 g, 0.33 mmol, 6 eq), AcOH (0.030 mL, 0.55 mmol, 10 eq), and piperidine (0.055 mL, 0.55 mmol, 10 eq). The reaction mixture was stirred o/n. The reaction was monitored by reverse phase TLC (Water/Acetonitrile 7:3). The reaction mixture was concentrated under vacuum and co-evaporated with toluene and the residue was purified by reverse flash column chromatography (Water/Acetonitrile 94:6 to 85:15 then to 80:20) to obtain **Red BODIpH 1** (5.8 mg, 10 % yield) as a purple solid and **fr BODIpH 1** (18.7 mg, 23 %) as blue solid with piperidinium as counterions.

*Red BODipH 1*

**$^1$H NMR** (300 MHz, Methanol-*d*$_4$) δ 8.52-8.51 (m, 1H), 8.38 (d, *J* = 16.8 Hz, 1H), 7.92 (d, J=1.2 Hz, 2H), 7.80(s, 1H), 7.76 (d, *J* = 16.8 Hz, 1H), 7.35 (dd, *J* = 8.4, 2.2 Hz, 1H), 7.10 (d, *J* = 8.4 Hz, 1H), 3.70 – 3.47 (m, 14H), 3.30-3.27 (m, 2H), 3.15 – 3.06 (m, 8H), 2.75 (q, J=7.2 Hz, 2H), 2.55 (s, 3H), 2-38 (q, J=7.53 Hz, 2H), 1.81 – 1.71 (m, 8H), 1.63 (m, 4H), 1.47 (d, *J* = 4.0 Hz, 6H), 1.19 (t, *J* = 7.4 Hz, 3H), 1.01 (t, *J* = 7.5 Hz, 3H). **$^{13}$C NMR** (75 MHz, Methanol-*d*$_4$) δ 168.97, 156.42, 148.15, 143.87, 143.07, 139.53, 139.45, 138.19, 137.24, 134.17, 133.76, 133.49, 132.58, 132.04, 131.64, 127.94, 127.42, 125.60, 125.14, 122.45, 118.51, 116.32, 70.22, 70.15, 69.99, 69.82, 69.62, 69.00, 50.32, 44.37, 39.19, 22.35,

21.66, 17.70, 16.46, 13.57, 11.60, 11.29, 10.78. **HRMS (ESI) m/z**: [M] $2^-$ Calculated for $C_{39}H_{45}BF_2N_6O_{11}S_2$ 443.1330. Found 443.1328.

*fr BODIpH 1*

**$^{1}$H NMR** (300 MHz, Methanol-$d_4$) δ 8.54-8.48 (m, 4H), 8.07-7.95 (m, 4H), 7.91 –7.79 (m, 3H), 7.43 (dd, *J* = 8.4 Hz, 2.1 Hz, 1H), 7.17 (d, *J* = 8.4 Hz, 1H), 3.74 – 3.48 (m, 14H), 3.31-3.27 (m, 2H), 3.18 – 3.07 (m, 16H), 2.80 (q, *J* = 7.4 Hz, 4H), 1.79 – 1.74 (m, 16H), 1.73 – 1.59 (m, 8H), 1.53 (s, 6H), 1.30 – 1.19 (m, 6H). **$^{13}$C NMR** (75 MHz, Methanol-$d_4$) δ 170.34, 162.97, 151.62, 145.54, 144.62, 140.82, 140.55, 138.30, 136.27, 134.95, 134.59, 129.51, 128.95, 127.34, 126.73, 126.47, 123.55, 120.12, 117.82, 71.57, 71.51, 71.34, 71.19, 70.97, 70.39, 51.68, 45.73, 40.55, 23.70, 23.02, 19.11, 14.86, 12.37. **HRMS (ESI) m/z:** [M] $4^-$ Calculated for $C_{46}H_{47}BF_2N_6O_{17}S_4$ 283.0491. Found 283.049.

**Red and fr BODIpH 2**

To a solution of **Yellow BODIpH 2** (0.046 g, 0.064 mmol, 1 eq) in DMSO (1 mL) under argon were 4-formylbenzene-1,3-disulfonate (0.119 g, 0.38 mmol, 6 eq), AcOH (0.040 mL, 0.64 mmol, 10 eq), and piperidine (0.065 mL, 0.64 mmol, 10 eq). The reaction mixture was heated to 25 °C o/n. The reaction was monitored by reverse phase TLC (Water/Acetonitrile 6:4). The reaction mixture was concentrated under vacuum and co-evaporated with toluene and the residue was purified by reverse flash column chromatography (Water/Acetonitrile 95:5 to 85:15 then to 80:20) to obtain **Red BODIpH 2** (1.8 mg, 3 % yield) as a purple solid and **fr BODIpH 2** (32 mg, 31%) as blue solid with piperidinium as counterions.

*Red BODIpH 2*

**$^{1}$H NMR** (300 MHz, Methanol-$d_4$) δ 8.54 – 8.52 (m, 1H), 8.39 (d, *J* = 16.8 Hz, 1H), 7.93 – 7.91 (m, 2H), 7.80 – 7.75 (m, 2H), 7.59 – 7.57 (m, 1H), 3.67 – 3.58 (m, 16H), 3.14 – 3.09 (m, 8H), 2.78 – 2.73 (m, 2H), 2.55 (s, 3H), 2.42-2.39 (m, 2H), 1.79 –1.75 (m, 8H), 1.69 – 1.67 (m, 4H), 1.58 – 1.55 (m, 6H), 1.28 – 1.19 (m, 3H), 1.05 – 1.01 (m, 3H). **$^{13}$C NMR** (75 MHz, Methanol-$d_4$) δ 170.52, 157.90, 149.49, 145.15, 144.37, 140.75, 139.45, 138.62, 137.27, 135.63, 135.16, 133.97, 133.45, 132.95, 128.81, 126.99, 126.49, 123.86, 71.54, 71.48, 71.34, 71.19, 70.97, 70.44, 51.68, 45.73, 40.60, 30.70, 23.71, 23.03, 19.08, 17.84, 14.93, 12.99, 12.88, 12.36. **HRMS (ESI) m/z**: [M] $2^-$ Calculated for $C_{39}H_{44}BBrF_2N_6O_{11}S_2$ 482.0882. Found 482.0882.

*fr BODIpH 2*

**$^{1}$H NMR** (300 MHz, Methanol-$d_4$) δ 8.58 – 8.45 (m, 4H), 8.05-7.97 (m, 4H), 7.87-7.82 (m, 3H), 7.63 (s, 1H), 3.74 – 3.54 (m, 16H), 3.11 (t, *J* = 5.5 Hz, 16H), 2.81 (q, *J* = 7.2 Hz, 4H), 1.80 – 1.72 (m, 16H), 1.68 – 1.65 (m, 8H), 1.61 (s, 6H), 1.25 (t, *J* = 7.4 Hz, 6H). **$^{13}$C NMR** (75 MHz, Methanol-$d_4$) δ 169.22, 158.83, 150.64, 144.35, 143.41, 139.23, 137.06, 136.87, 136.77, 135.16, 133.55, 133.47, 127.56, 126.55, 126.41, 126.00, 125.11, 122.10, 116.26, 112.18, 70.23, 70.14, 69.96, 69.76, 69.61, 68.73, 50.32, 48.49, 48.21, 47.93, 47.64, 47.36, 47.08, 46.79, 44.38, 39.44, 22.33, 21.66, 17.75, 13.49, 11.20. **HRMS (ESI) m/z:** [M] $4^-$ Calculated for $C_{46}H_{47}BBrF_2N_6O_{17}S_4$ 403.7047. Found 403.7045.

**LipH**

DBCO-PEG (5000)- DSPE (1.5 mg, 0.25 µmol) was added to a solution of **fr BodipH 2** (0.75 mg, 0.5 µmol) in DMF under argon. The reaction mixture was stirred overnight. The reaction was monitored by reverse phase TLC (Water/Acetonitrile 1:9). The reaction mixture was concentrated under vacuum and the residue was purified by steric exclusion chromatography column LH20 stationary phase (DCM/MeOH (1:1)) to obtain LipH as a blue solid (2 mg, quant).

#### V. NMR spectra

*Compound 2*

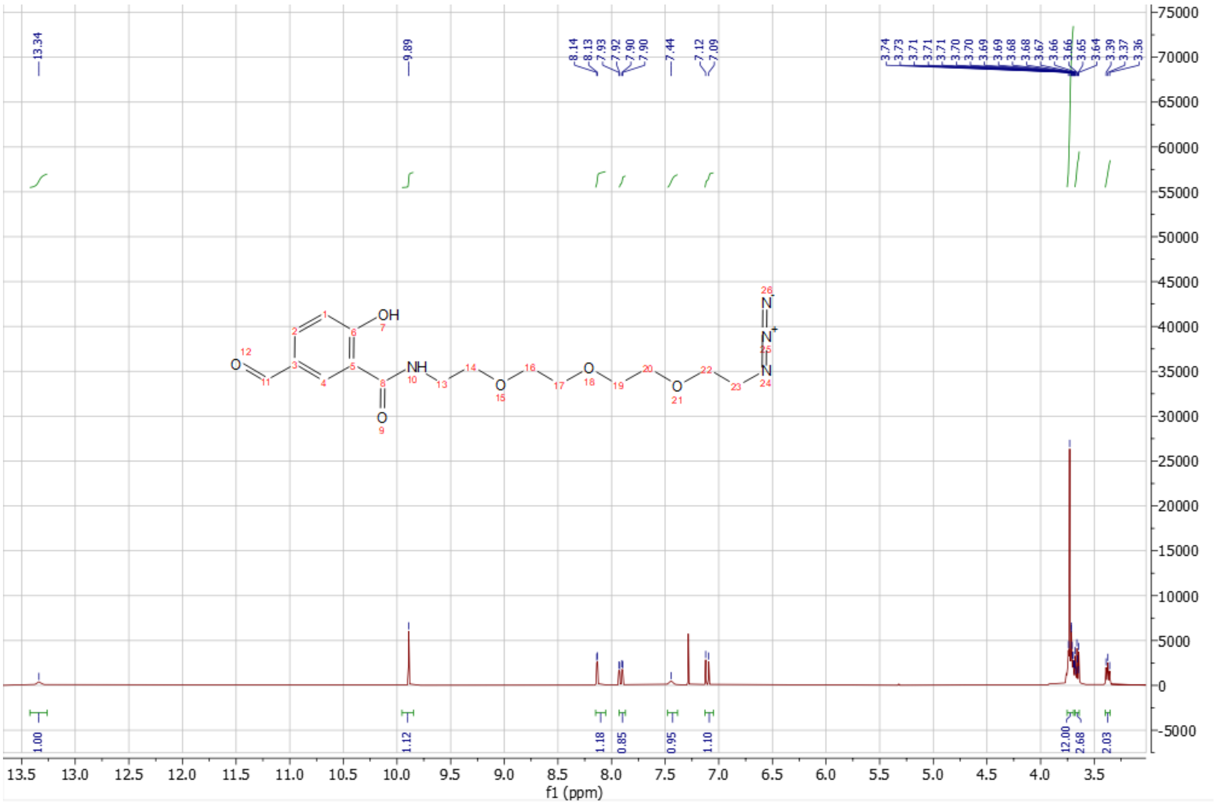

r

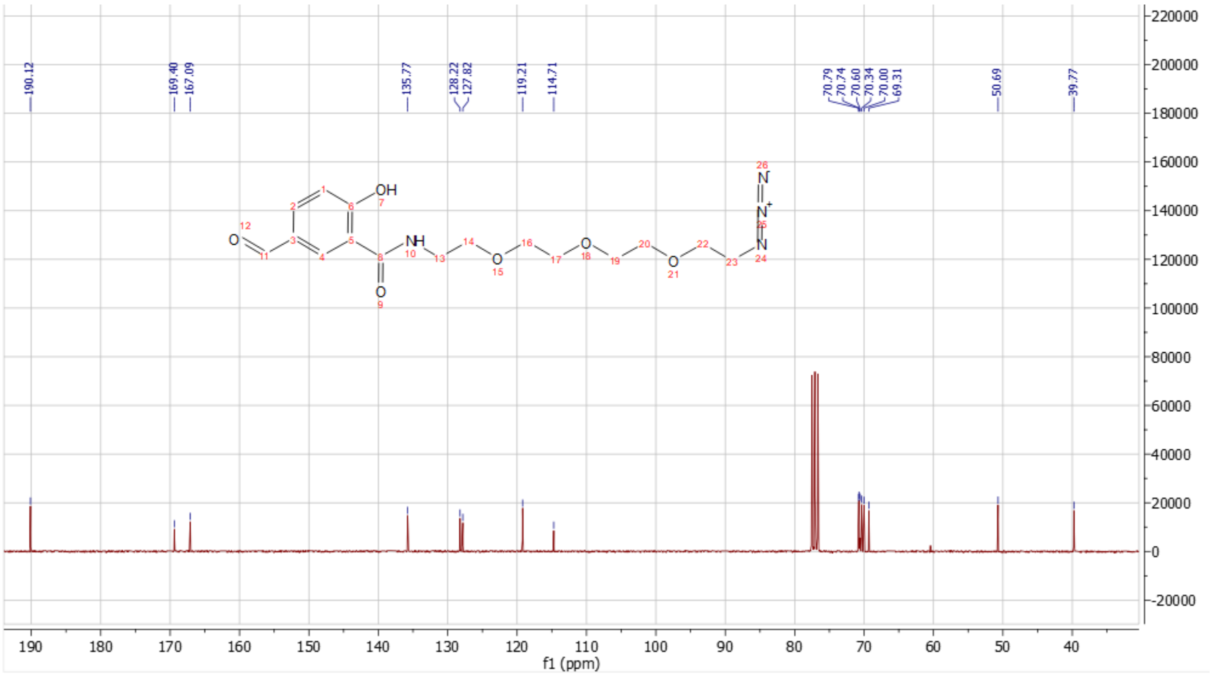

*Compound 3*

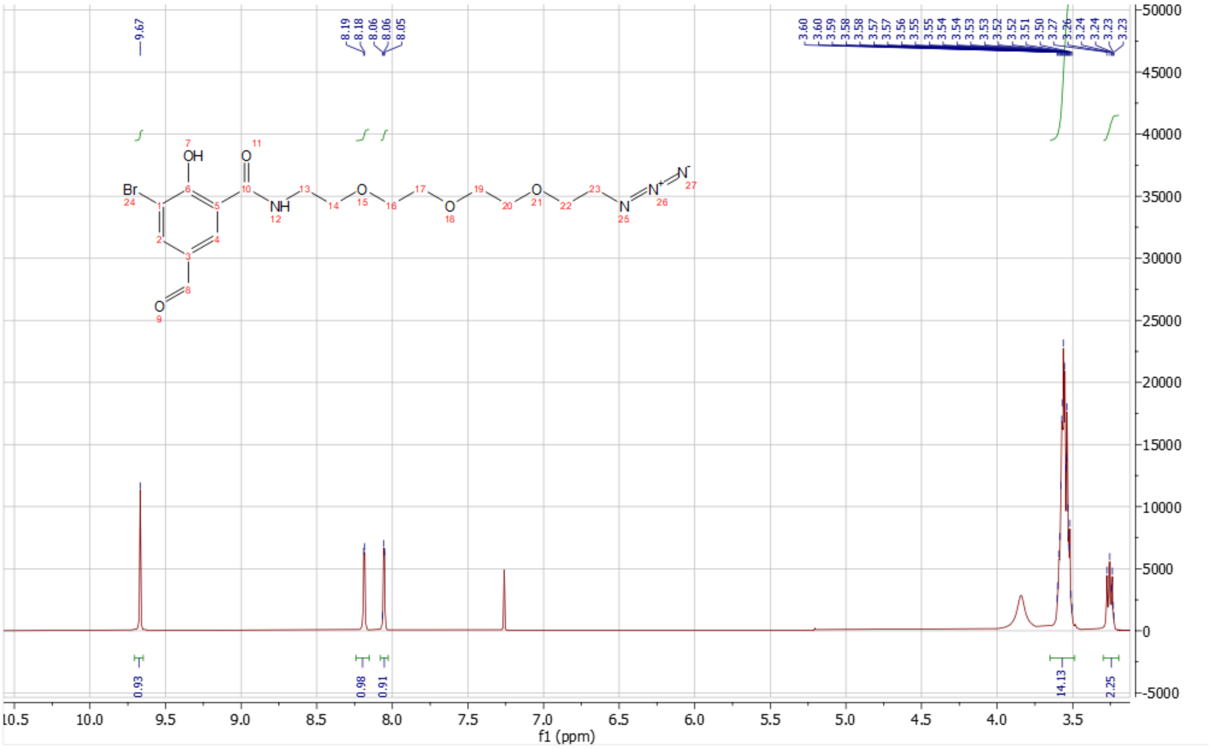
9.67
8.19
8.18
8.06
8.06
8.05
3.60
3.60
3.59
3.58
3.58
3.57
3.57
3.56
3.55
3.55
3.54
3.54
3.53
3.53
3.52
3.52
3.51
3.50
3.27
3.26
3.24
3.24
3.23
3.23
0.93
0.98
0.91
14.13
2.25
f1 (ppm)

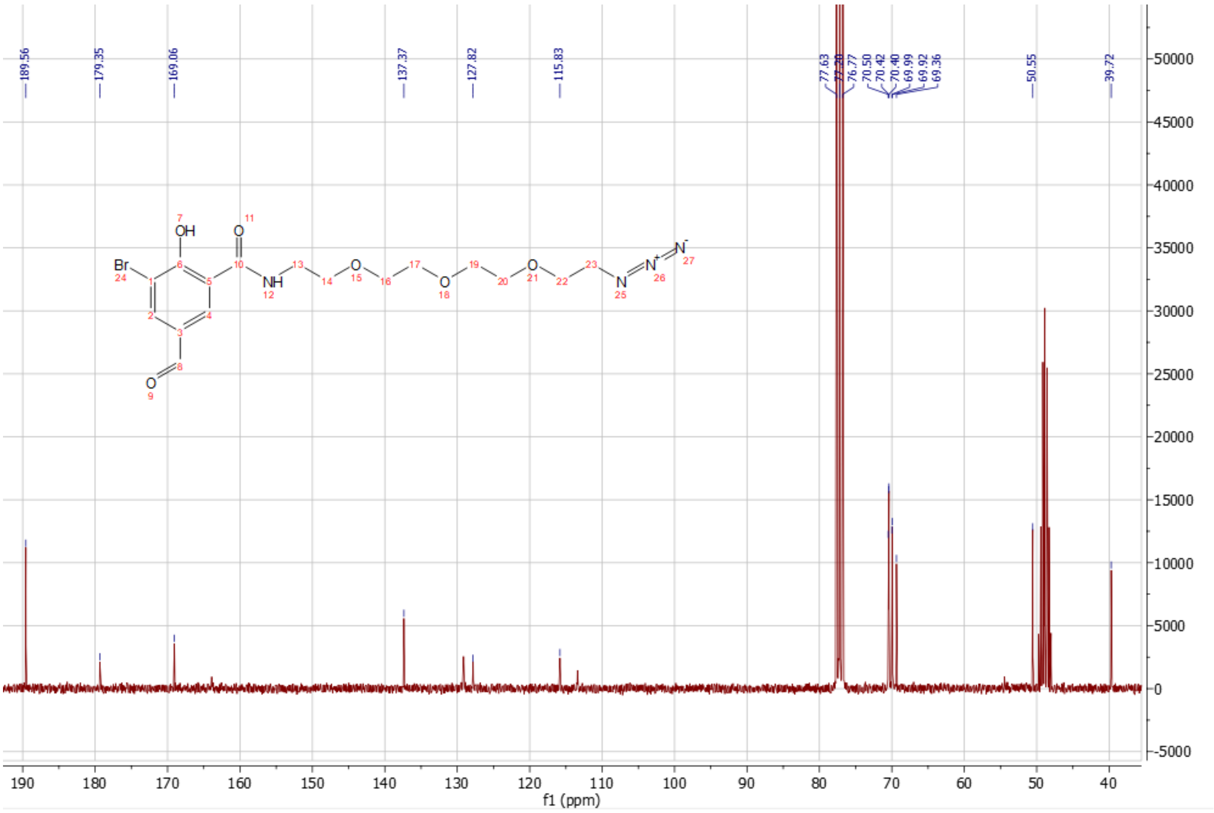
189.56
179.35
169.06
137.37
127.82
115.83
77.63
77.20
76.77
70.50
70.42
70.40
69.99
69.92
69.36
50.55
39.72
f1 (ppm)

*Yellow BODIpH 1*

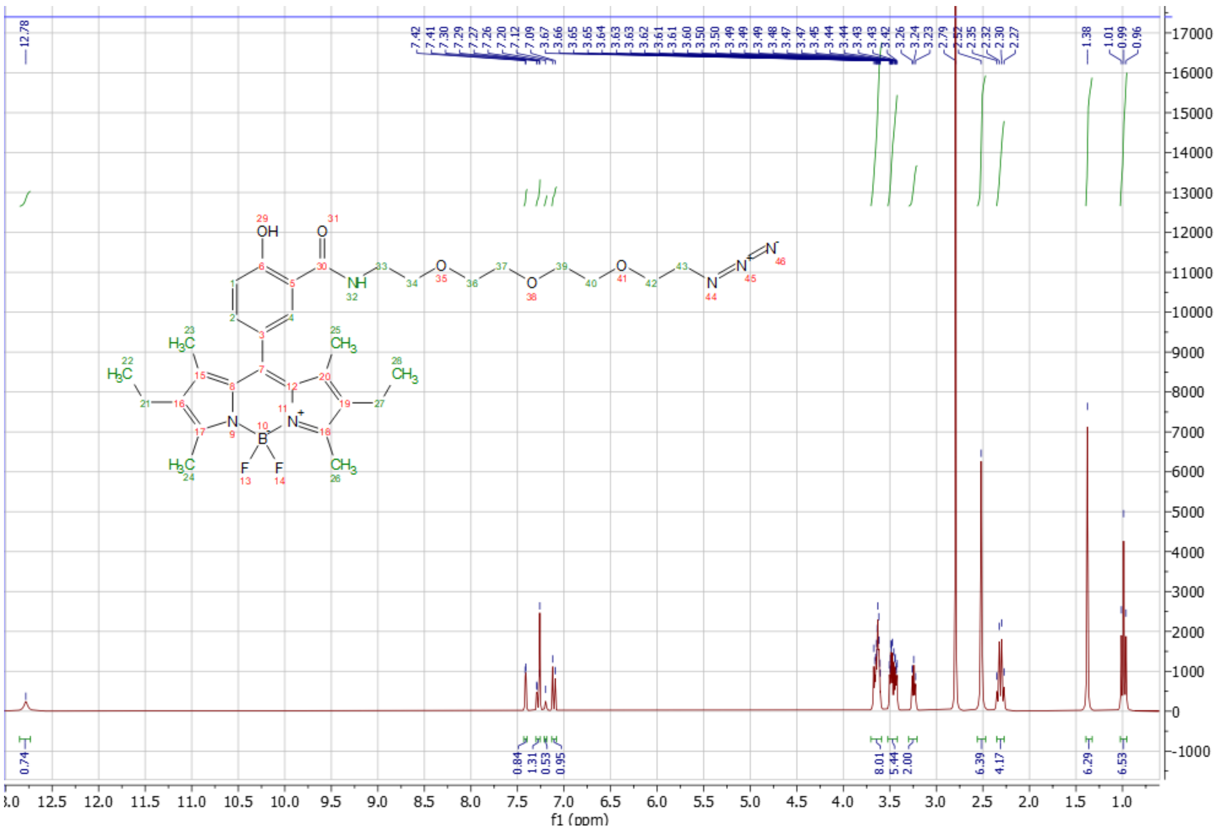

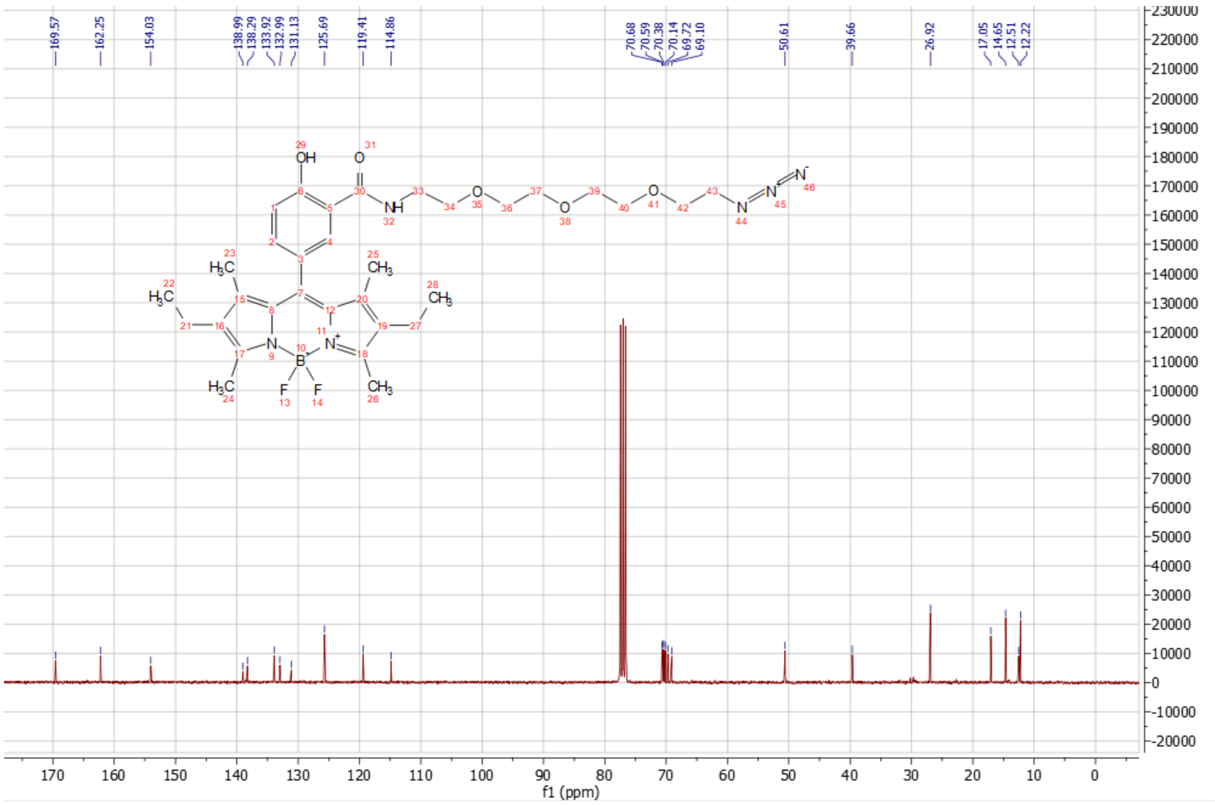

*Yellow BODIpH 2*

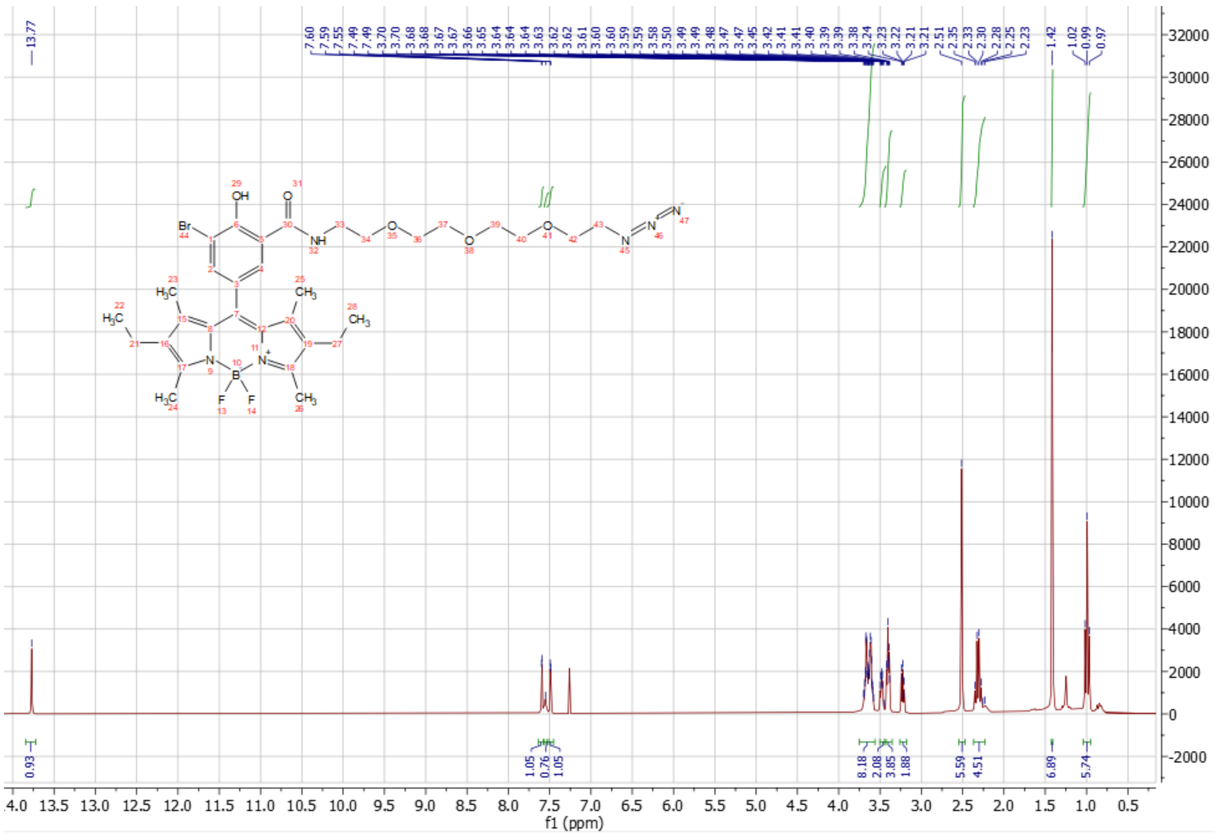

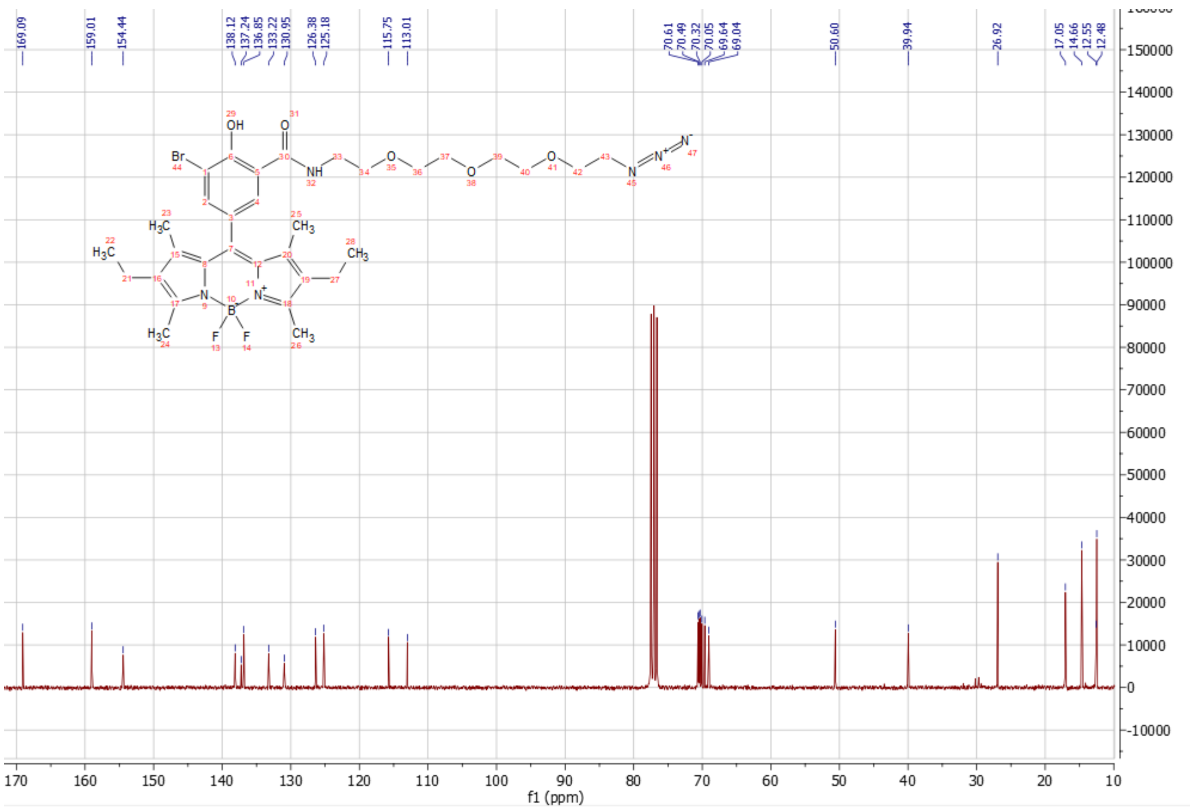

*Red BODIpH 1*

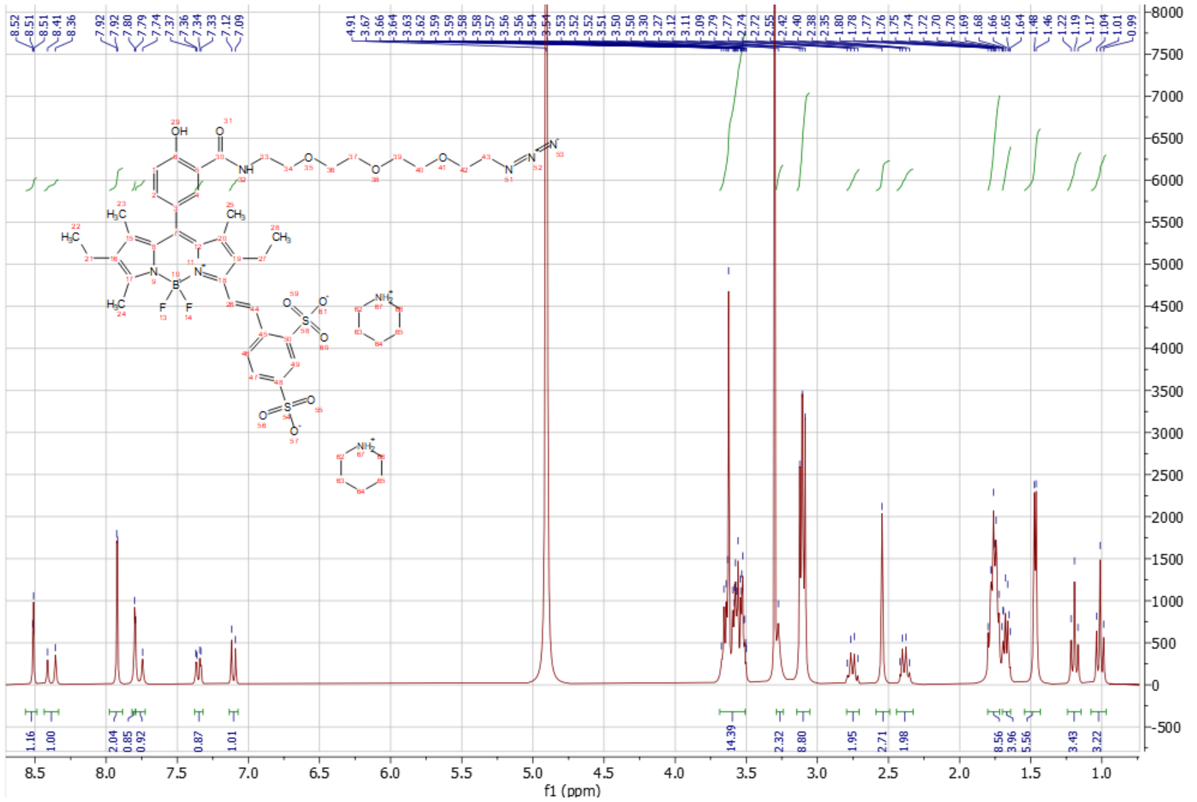

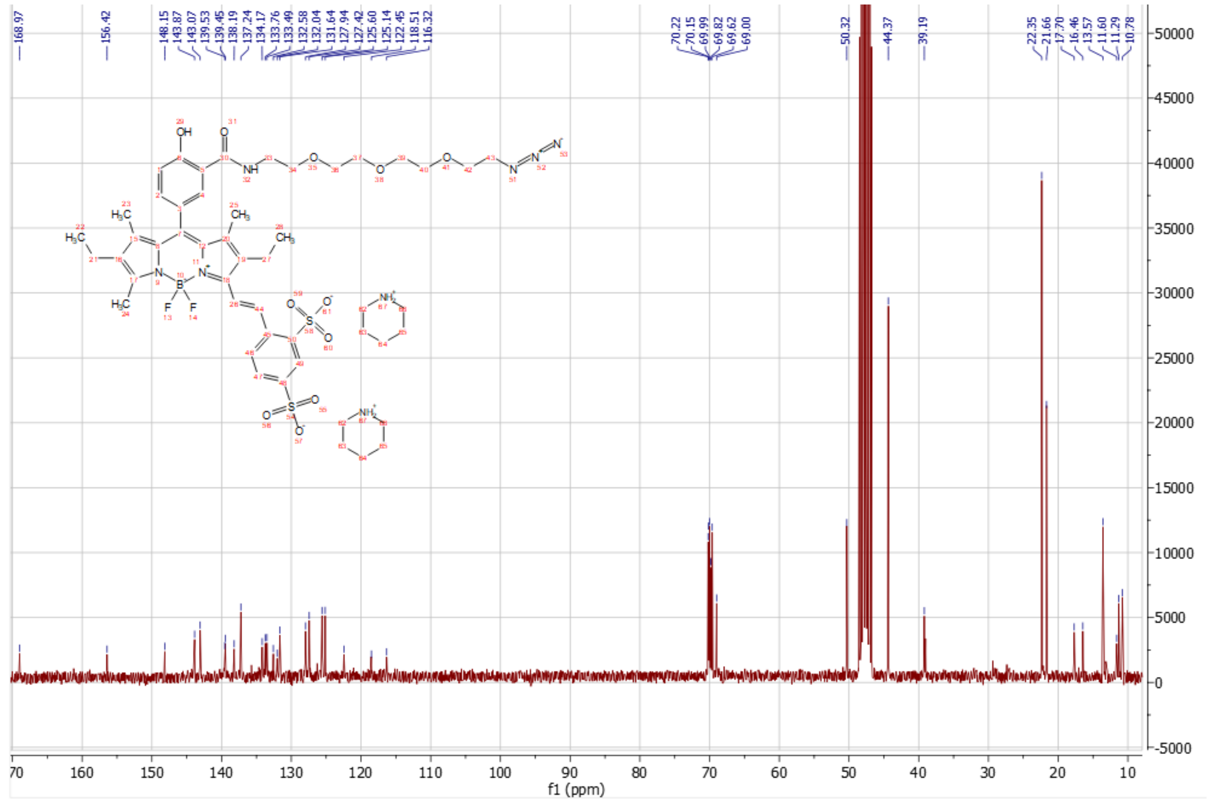

*fr BODIpH 1*

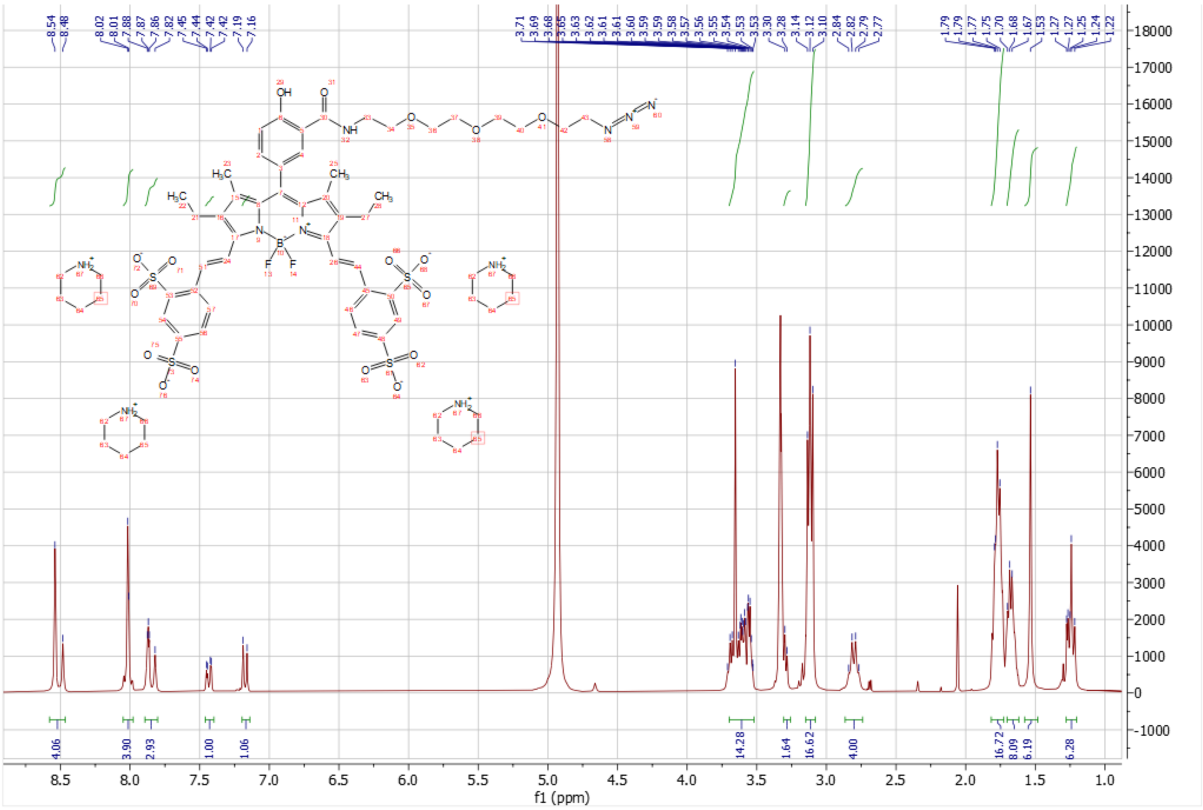

8.5
8.0
7.5
7.0
6.5
6.0
5.5
5.0
4.5
4.0
3.5
3.0
2.5
2.0
1.5
1.0
f1 (ppm)
4.06
3.90
2.93
1.00
1.06
14.28
1.64
16.62
4.00
16.72
8.09
6.19
6.28

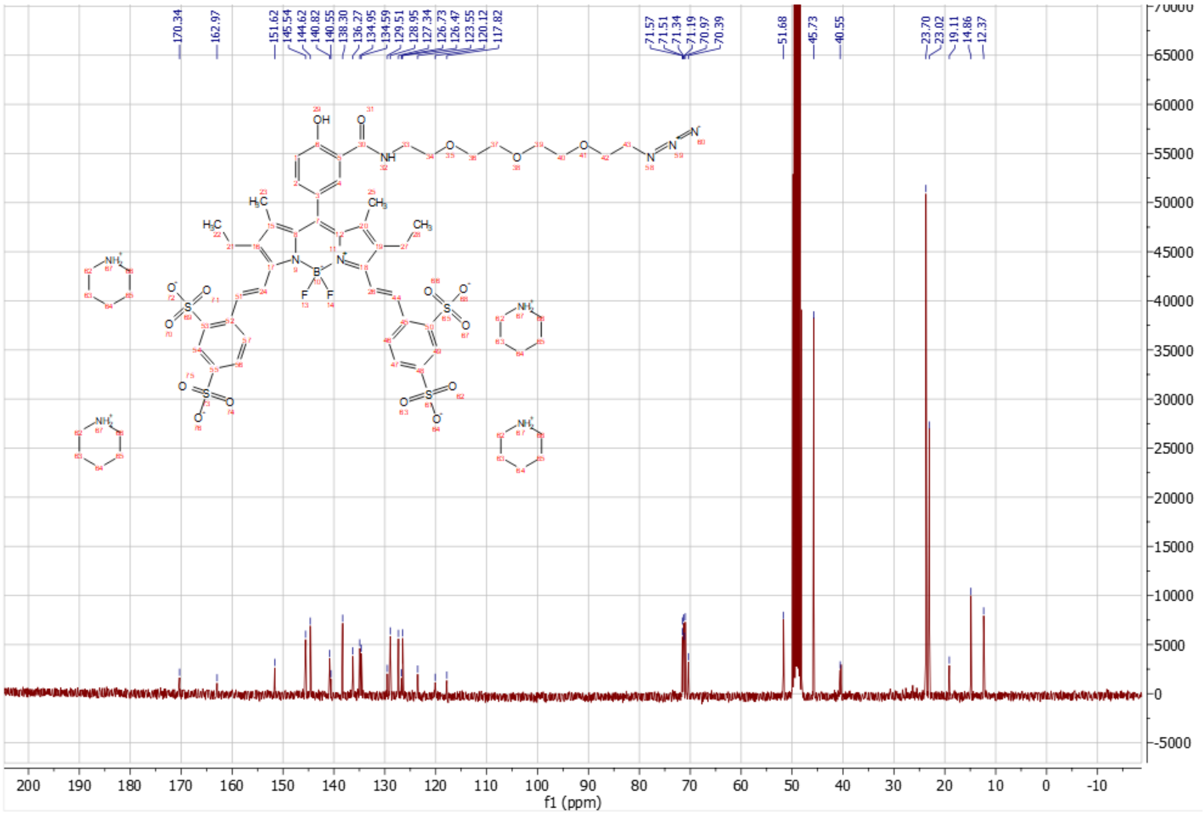

*Red BODIpH 2*

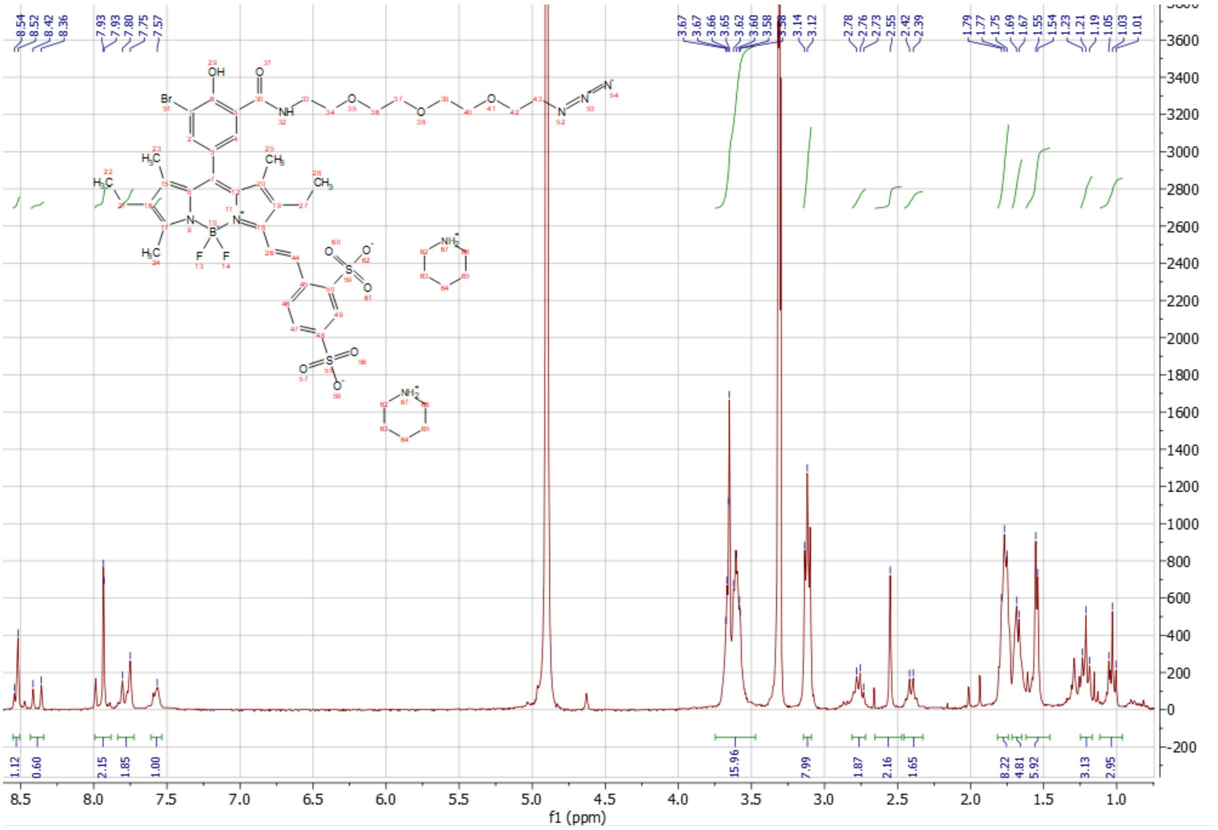

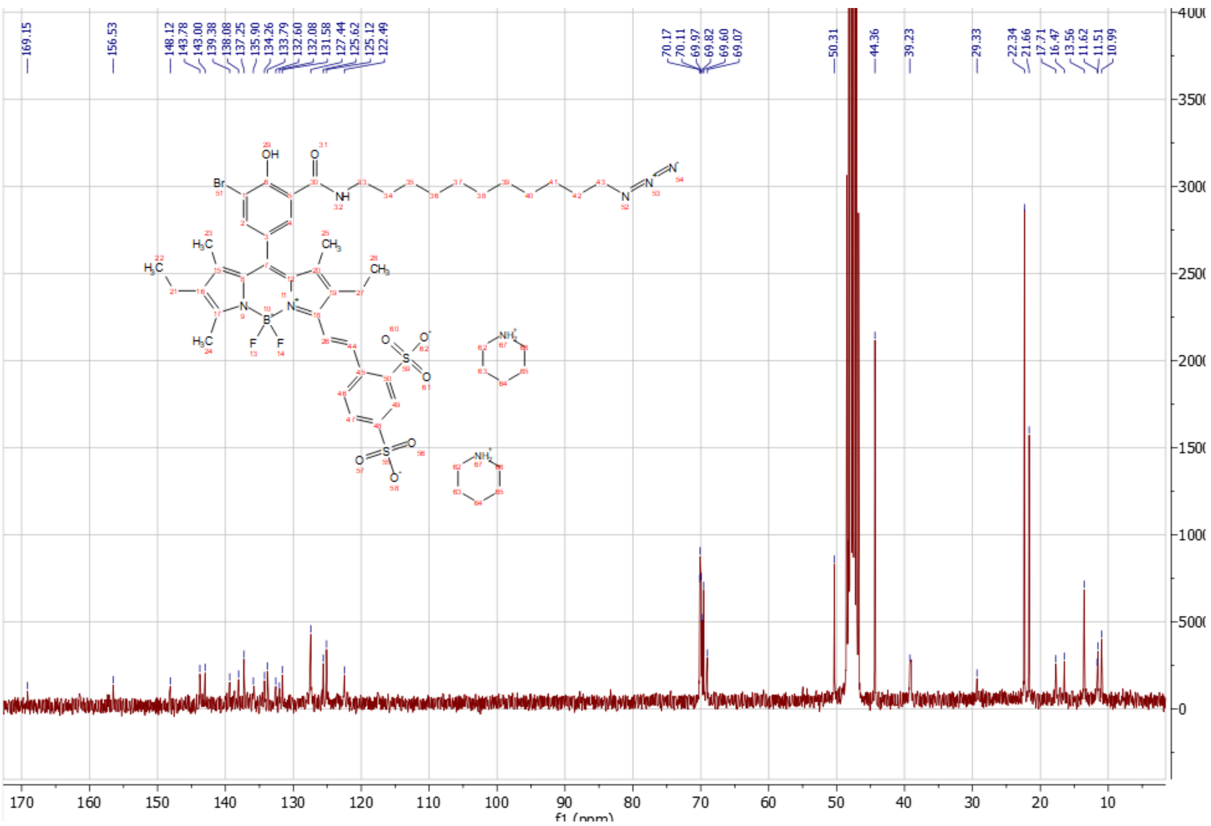
169.15
156.53
148.12
143.78
143.00
139.38
138.08
137.25
135.90
134.26
133.79
132.60
132.08
131.58
127.44
125.62
125.12
122.49
70.17
70.11
69.97
69.82
69.60
69.07
50.31
44.36
39.23
29.33
22.34
21.66
17.71
16.47
13.56
11.62
11.51
10.99
f1 (ppm)

*fr BODIpH 2*

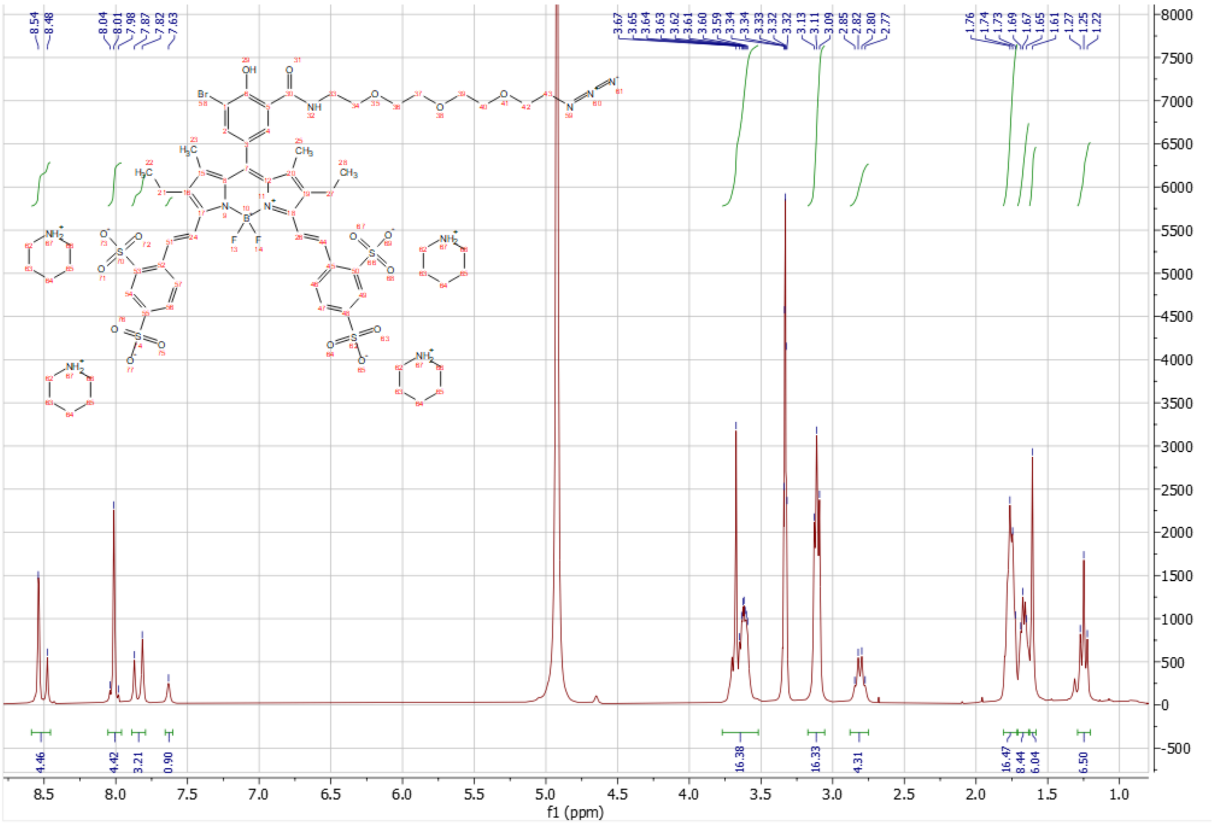
8.54 8.48
8.04 8.01 7.98 7.87 7.82 7.63
3.67 3.65 3.64 3.63 3.62 3.61 3.60 3.59 3.34 3.34 3.33 3.32 3.32 3.13 3.11 3.09 2.85 2.82 2.80 2.77
1.76 1.74 1.73 1.69 1.67 1.65 1.61 1.27 1.25 1.22
4.46 4.42 3.21 0.90 16.38 16.33 4.31 16.47 8.44 6.04 6.50
f1 (ppm)

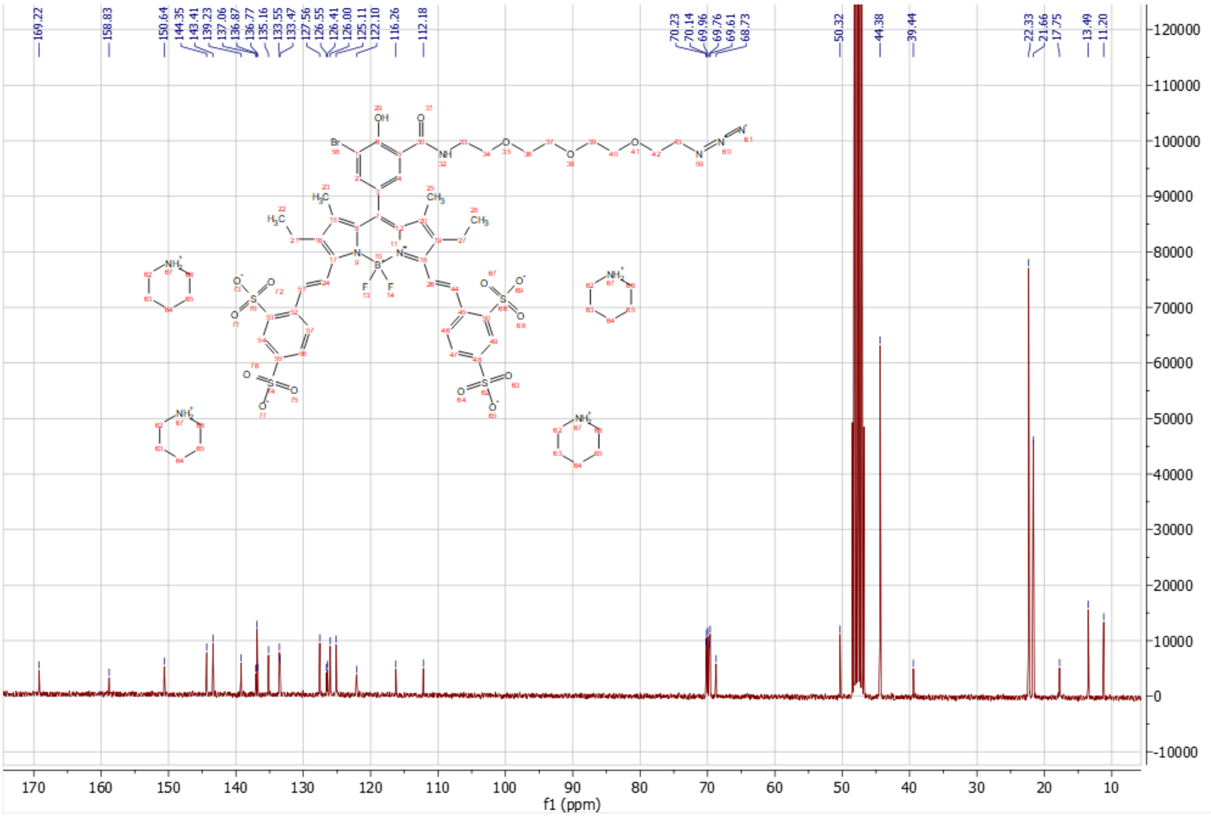
169.22
158.83
150.64 144.35 143.41 139.23 137.06 136.87 136.77 135.16 133.55 133.47 127.56 126.55 126.41 126.00 125.11 122.10
116.26
112.18
70.23 70.14 69.96 69.76 69.61 68.73
50.32
44.38
39.44
22.33 21.66 17.75
13.49 11.20
f1 (ppm)
